# Direct energy transfer from photosystem II to photosystem I is the major regulator of winter sustainability of Scots pine

**DOI:** 10.1101/2020.06.10.144170

**Authors:** Pushan Bag, Volha Chukhutsina, Zishan Zhang, Suman Paul, Alexander G. Ivanov, Tatyana Shutova, Roberta Croce, Alfred R. Holzwarth, Stefan Jansson

## Abstract

Evergreen conifers in boreal forests can survive extremely cold (freezing) temperatures during the long dark winter and fully recover during the summer. A phenomenon called ‘sustained quenching’ putatively provides photoprotection and enables their survival, but its precise molecular and physiological mechanisms are not understood. To unveil them, we have analyzed the seasonal adaptation of the photosynthetic machinery of Scots pine (*Pinus sylvestris*) trees by monitoring multi-year changes in weather, chlorophyll fluorescence, chloroplast ultrastructure, and changes in pigment-protein composition. Recorded Photosystem II and Photosystem I performance parameters indicate that highly dynamic structural and functional seasonal rearrangements of the photosynthetic apparatus occur. Although several mechanisms might contribute to ‘sustained quenching’ of winter/early spring pine needles, time-resolved fluorescence analysis shows that extreme down-regulation of photosystem II activity along with direct energy transfer from photosystem II to photosystem I plays a major role. This mechanism is enabled by extensive thylakoid destacking allowing for mixing of PSII with PSI complexes. These two linked phenomena play crucial roles in winter acclimation and protection.

**Graphical abstract:** 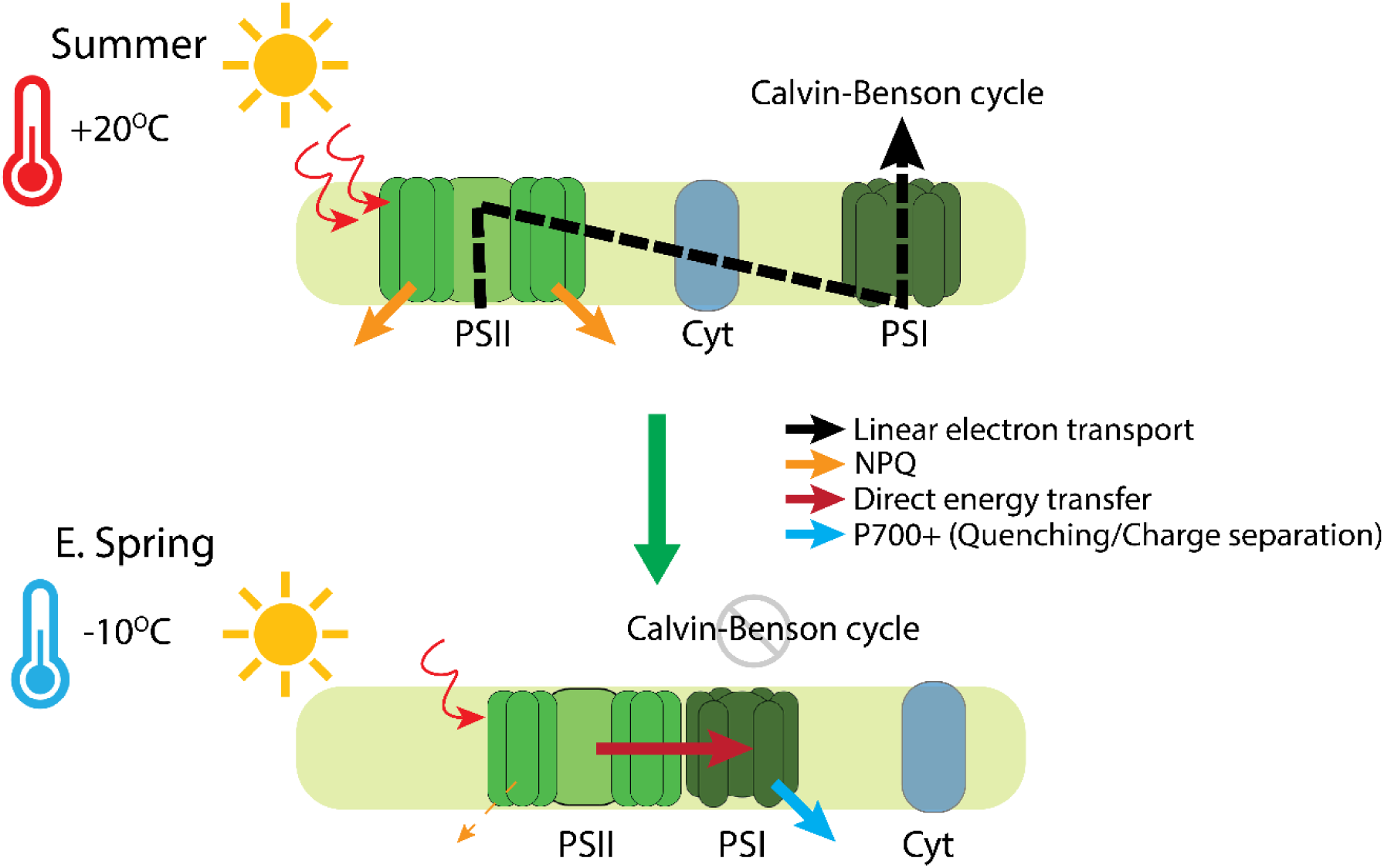

## Introduction

Photosynthesis is the basis for most life on earth, and the ability to sustain growth by harvesting sunlight confers such an enormous evolutionary advantage that photosynthetic organisms have developed numerous adaptations that enable them to photosynthesize in diverse environments. These include vast boreal forests, which cover ∼70% of all coniferous forest of the world (mainly in the northern hemisphere)^1^, which have lower species diversity than many other terrestrial ecosystems and are often dominated by evergreen conifers, like pine and spruce^2^. Broadleaf deciduous trees and shrubs are also present, and sometimes deciduous conifers such as Larch (*Larix*) species, but there is no doubt that evergreen conifers are particularly well adapted to harsh boreal conditions, including short growing seasons and cold, snowy winters. Thus, evergreen conifers’ adaptations to boreal regions must presumably include regulatory processes that protect the photosynthetic apparatus in their needles during the harsh winter and early spring. Knowledge of photoprotective mechanisms in various kinds of photosynthetic organisms has increased considerably in recent decades. Several control systems have been identified that allow the photosynthesis machinery to harmlessly dissipate excess excitation energy^3–7^. In 2003, Öquist and Huner^8^ published a seminal review of photosynthesis in evergreen plants, which pointed out that overwintering conifer needles enter a state of ‘sustained quenching’ during winter. They found strong evidence for major alterations in the organization and composition of the photosystem II (PS II) antenna but also concluded that photosystem I (PS I) may play an important role, via nonphotochemical quenching of absorbed light or via quenching absorbed light photochemically through cyclic electron transport. Recently it was also shown that alternative electron transport might add up to this as well^9^. Up to date there exists no clear mechanism rationalizing winter quenching in conifers, although there are several proposed hypotheses^8,10^. Several protection mechanisms may also occur in parallel, and (if so) they are likely activated largely well before the extreme stress occurs^11,12^. But the major conundrum arises in early spring, when temperatures are typically still low, so plants’ biochemical and metabolic activities are strongly limited, but solar radiation is already high^8^. Once induced, the mechanisms need to be sustained, i.e. locked-in over the long winter season and protect the photosynthetic machinery. When conditions improve late in the spring, photosynthetic apparatus is restored and attains its active growth state in summer.

In previous explorations of these phenomena, a marked drop in steady-state fluorescence (measured as Fv/Fm) has been recorded in several overwintering conifers, and some indications of the quenching mechanisms involved have been obtained. These reportedly include destacking of thylakoid membranes and associated changes in chloroplast ultrastructure ^13^, in accordance with the drop in PSII maximal fluorescence ^8^. In higher plants, thylakoid stacking and heterogeneity play crucial roles in the localizations of PSII and PSI in grana (tightly appressed thylakoid layers) and stroma lamellae, respectively. In conifers, although an extreme drop in maximal fluorescence in winter/early spring has been observed, its mechanistic relationship to thylakoid structural changes has not been explained. In the study reported here, we monitored steady-state chlorophyll fluorescence, ultrafast time-resolved fluorescence, and chloroplast ultrastructure in Scots pine needles from autumn to summer in three successive years. Our data strongly indicate that chlorophyll fluorescence quenching and thylakoid destacking, are strongly linked, mutually dependent, and crucial for the survival of evergreen conifers in the extreme northern boreal winter and early spring, when temperatures are low but solar radiation levels may be high.

## Results

### Seasonal changes in Photosystem II- and Photosystem I-related functional activities

To monitor photosynthetic performance, we recorded several PSII and PSI parameters during three consecutive growth seasons (2015-2016, 2016-2017, and 2017-2018) along with concomitant changes in daily air temperature and solar radiation (Fig. 1A-I, II, III). For simplicity, in the main figures we only present (here onwards) data from 2017-2018, which we divided into five distinct seasons, based on weather parameters: Summer (S, June-Aug), autumn (A, Sept-mid Nov), winter (W, mid Nov-mid Feb), early spring (ES, mid Feb-mid Apr), and late spring (LS, mid Apr-June). Data from the other two growth seasons are provided in Supplementary information 1, 2 and 3.

**Figure 1.**
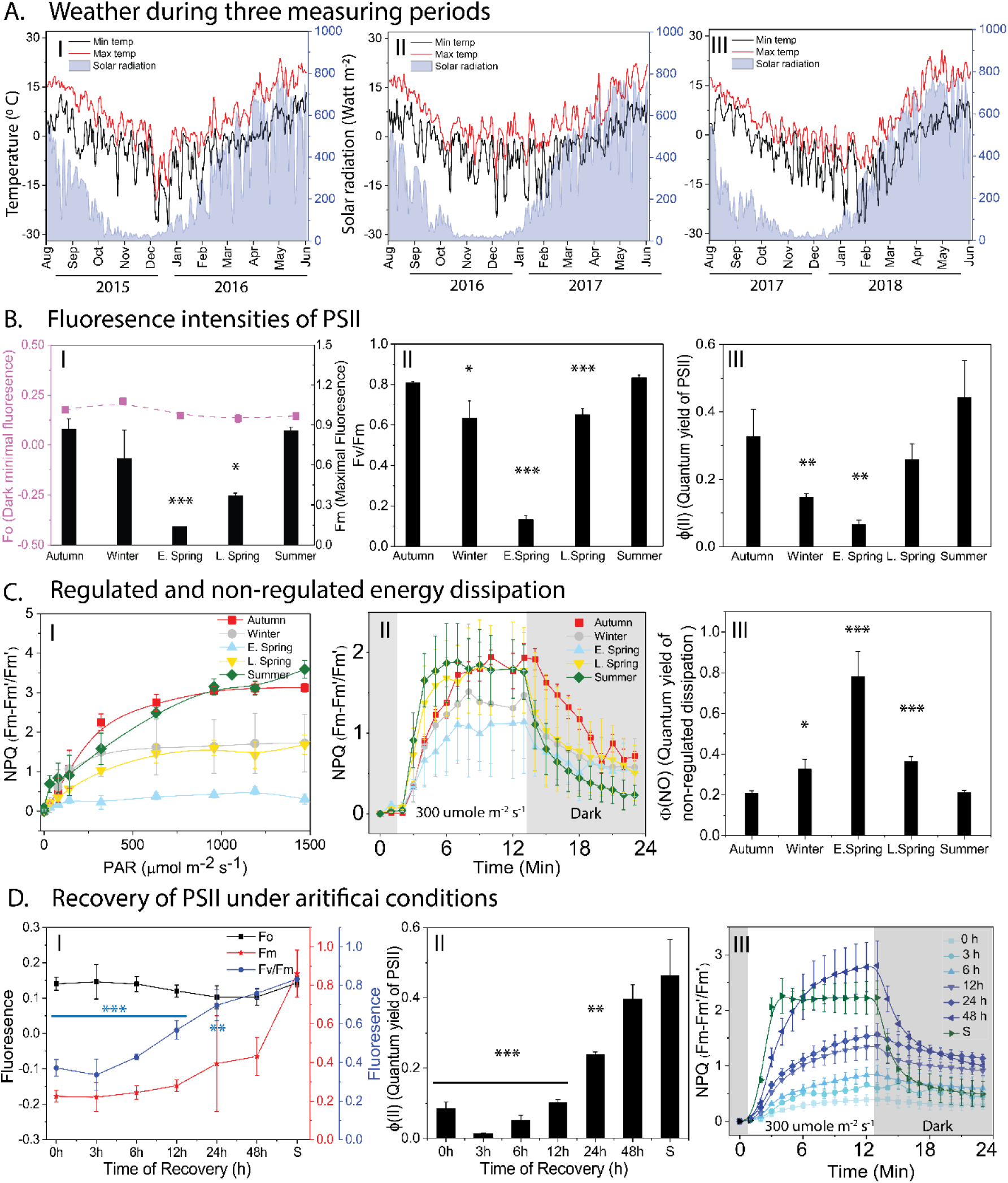
Seasonal dynamics of weather and photochemical performance of PSII measured by chlorophyll fluorescence in Scots pine needles. A. Changes in temperature (°C) (Left Y axis) and solar radiation (watt m-2) (Right Y axis) during 2015-2016 (I), 2016-2017 (II), 2017-2018 (III) measuring seasons. B. Seasonal dynamics of PSII photochemistry: (I) Changes in maximal (Fm) and basic (Fo) fluorescence. (II) Maximal quantum efficiency of PSII measured as Fv/Fm (III) Effective quantum yield of PSII (Φ(II)). C. Energy dissipation measured as regulated and non-regulated non photochemical quenching. (I) Changes in NPQ with increasing PAR (II) Induction of NPQ with constant actinic light, (III) Quantum yield of non-regulated non photochemical quenching. D. Recovery of pine needles under artificial conditions at 80 μmol photons m^-2^ s^-1^of light for 48h with 18/6 photoperiod (I) Changes in Fo, Fm and Fv/Fm, (II) Changes in Φ(II), (III) Induction of NPQ with constant actinic light of 300 μmol photons m^-2^ s^-1^. All measurements were taken after 30 mins of dark adaptation at 4°C in winter and room temperature in summer. All data are means ± SD (n = 3-5) and the statistically significant differences are marked by asterisks: * - p ≤ 95%; ** - p ≤ 99%.; *** - p ≤ 99.9%.

We found characteristic seasonal patterns in maximum PSII fluorescence (Fm) and maximum quantum efficiency of PSII (expressed as Fv/Fm), in accordance with earlier reports ^8,10^. Fv/Fm was highest in S, fell with reductions in ambient temperatures during A and W (Fig. 1B-I, II), and was lowest in ES (63% lower than in S), when low temperatures coincided with rises in solar irradiance (Fig. 1A-I-III; see also SI 2A). During LS, Fv/Fm gradually increased with increasing temperatures and peaked in S. The seasonal changes recorded in Fv/Fm were found to be mostly affected by changes in Fm rather than Fo (basic intrinsic fluorescence) (Fig. 1B-I). For deeper understanding of PSII performance, the fraction of absorbed light energy utilized by PSII photochemistry^14^ (ΦPSII) was measured (Fig. 1B-III). During W ΦPSII decreased significantly and reached minimal values (19% of S values) in ES (SI 2B).

Furthermore we tried to quantify the amount of absorbed light energy thermally dissipated by non-photochemical quenching (NPQ)^3^. The component of NPQ that play a crucial role under fluctuating light conditions is the fast component (ΔpH-PsbS dependent-qE^3^ or zeaxanthin dependent-qZ^4^), which increased with increasing light intensities in all the seasons. In S and A, the fast component did not reach a stationary phase even at 1500 μmol m^-2^ sec^-1^ illumination, while in W and LS samples a stationary phase was reached at 500 μmol m^-2^ sec^- 1^ and in ES already at 300 μmol m^-2^ sec^-1^ (Fig. 1C-I). Most importantly, in ES inducible steady-state NPQ was much smaller and overall slower due to the smaller amplitude of the fast component (∼50% less than in S) in ES (Fig. 1C-II). This is due to the fact that ES needles have already developed static NPQ. Instead the quantum yield of non-regulated and/or constitutive loss (ΦNO) of energy was high during ES (Fig. 1C-III) (SI 2D). This strongly suggests that this static NPQ is the fraction of absorbed light energy neither going to drive photochemistry (Φ_PSII_) nor thermally dissipated by rapid regulated NPQ processes (qE/qZ).

To confirm that this static quenching is a manifestation of a ‘sustained mode of quenching’ we artificially relaxed the ES needles (hereafter, ESR) for 48h in low light (80 μmol m^-2^ sec^-1^) with 18/6 h photoperiod. During this recovery we observed no significant changes in Fv/Fm for 6-8 hours (less than a 10% increase), a modest increase (30%) after 24 h (Fig. 1D-I) and almost complete (95%) recovery to S levels after 48 h. Fo did not change significantly throughout the recovery period (Fig. 1D-I), but changes in Fm followed the same recovery dynamics as Fv/Fm (Fig. 1D-I), although maximal fluorescence (Fm) was much higher in S than in 48 h ESR samples. ΦPSII did not change significantly until 12 h of recovery but recovered to 50% and 90% of S levels after 24 and 48 h, respectively (Fig. 1D-II). qE/qZ-dependent fast NPQ kinetics followed similar patterns to ΦPSII (Fig. 1D-III). It should be noted that light-dependent induction of the fast component of NPQ (qE/qZ) was slower in the ESR samples than in S samples, even after 48 h of recovery, which appears to be mainly due to photoinhibition.

Seasonal changes in partitioning of absorbed light energy within PSI showed that Y(I) was highest in S, declined in A and W, and reached minimal values in ES (Fig. 2A-I). Although it decreased, Y(I) was much less affected than the PSII activity (ΦPSII) during the cold periods (Fig. 1B-III) (SI 3A). Y(NA) followed the same seasonal pattern, with minimum values registered in ES and a steady recovery until S (Fig. 2A-II) (SI 3B). In contrast to Y(NA), Y(ND) considerably increased during the cold periods and peaked (at 69% higher than S values) in ES, then gradually declined to minimum values in S (Fig. 2A-III, SI 3C). This suggests that during early spring most of the absorbed light energy is dissipated by oxidized P700 (P700+) to prevent photoinhibition caused by over reduction of the acceptor side as iron sulfur clusters get damaged^15^. During the first 3 h of the recovery period, ΦPSI did not significantly change, but it gradually returned to S levels over 48 h (Fig. 2B-I). Y(NA) did not change much during recovery (Fig. 1E-II). Y(ND) increased slightly during the first 3 h, but then gradually decreased, and ESR samples did not quite reach S values. Based on these results, we conclude that the restoration of PSI light use efficiency takes longer than 48 hours.

**Figure 2.**
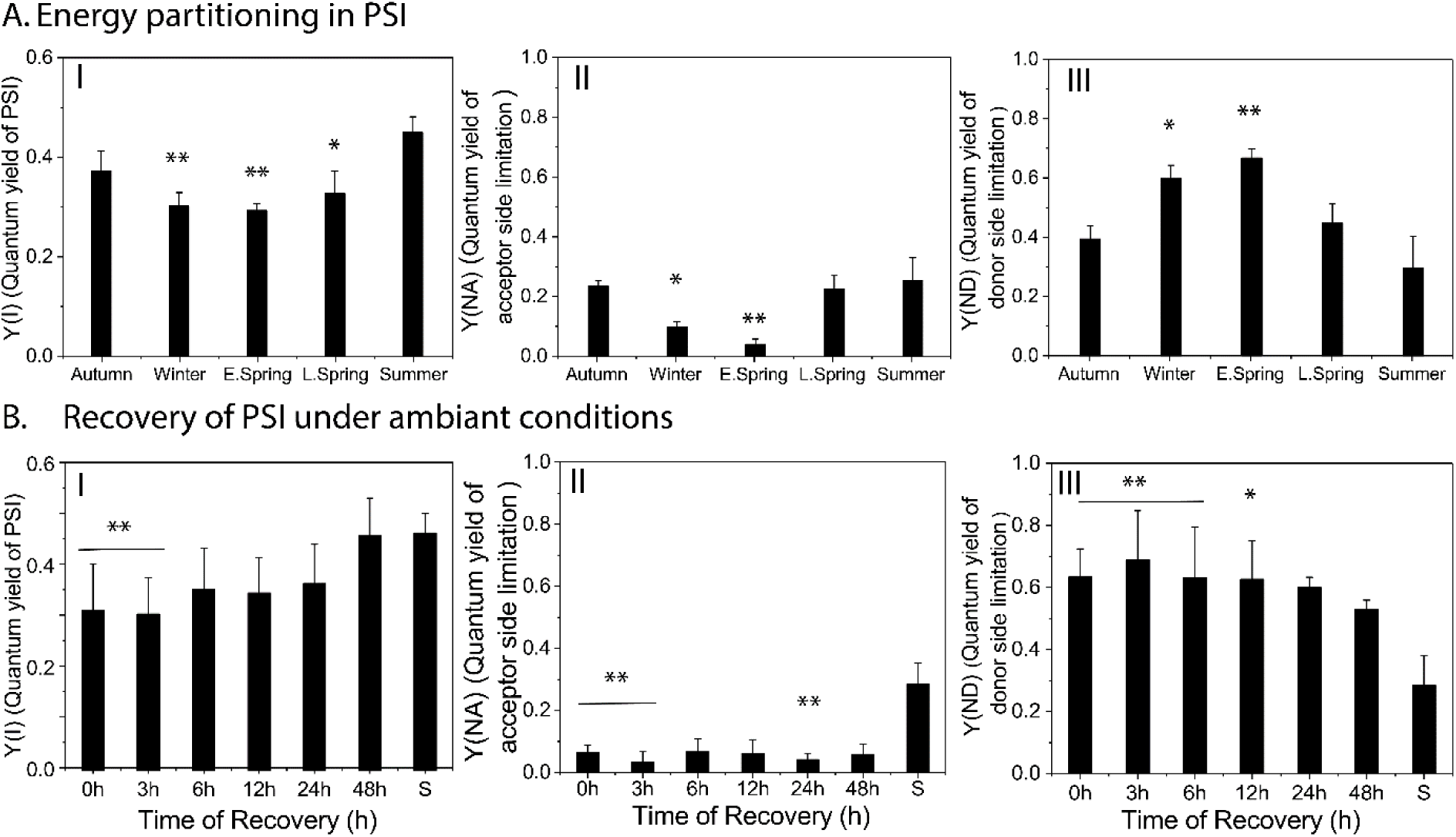
Seasonal changes of PSI photochemistry in Scots pine needles. **A**. Energy distribution in PSI considering Y(I)+Y(ND)+Y(NA) = 1, where Y(I) (I), Y(NA) (II) and Y(ND) (III) are photochemical quantum yield of PSI (when P700 is reduced and A is oxidised), energy dissipation in PSI (measure of acceptor side limitation, when P700 and A both are reduced) and energy dissipation in PSI (measure of donor side limitation, when P700 and A both are oxidised), respectively. **B**. Recovery of PSI photochemistry under ambient conditions: (I) Y(I), (II) Y(NA), (III) Y(ND) of PSI during the recovery period. All measurements were taken after 30 mins of dark adaptation at 4°C in winter and room temperature in summer. All data are means ± SD (n = 3-5) and the statistically significant differences are marked by asterisks: * - p ≤95%; ** - p ≤99%.

### Sustained non-photochemical quenching dominates in early spring needles

To elucidate the mechanism of “sustained quenching” observed in pine during the early spring periods, ultrafast time-resolved fluorescence measurements on intact pine needles were performed at a temperature of −20°C using the setup and approach described previously^16,17^. In addition, a special chamber was designed to keep the rotational cuvette with the sample at −20 C (SI 4.1). Fluorescence decay traces of S needles in the original state (dark-adapted, Fig. 3A, Summer dark) were compared to the needles collected in ES when sustained quenching was present (Fig. 3A, Early spring) and after the quenching in the needles had been relaxed at room temperature (ESR) (Fig. 3A, E. spring recovered). To understand whether sustained quenching observed in ES occurs via a similar quenching mechanism to light-induced quenching, S needles exposed to HL for 30 min (SQ) were also measured (Fig. 3A, Summer quenched). As shown by the direct comparison of the decay curves (Fig. 3A), the fluorescence in SQ needles is much shorter-lived as compared to that of S needles. However, the fluorescence decays of ES needles are still pronouncedly shorter-lived than that of the SQ samples, thus characterizing the “sustained quenching” in the ES samples as the most pronounced quenching at all detection wavelengths and under all conditions. In contrast, ESR and S samples showed very similar fluorescence decays, indicating that the ES sample recovered quite (although not completely, vide infra) well from sustained quenching within 48 h of recovery treatment.

**Fig 3.**
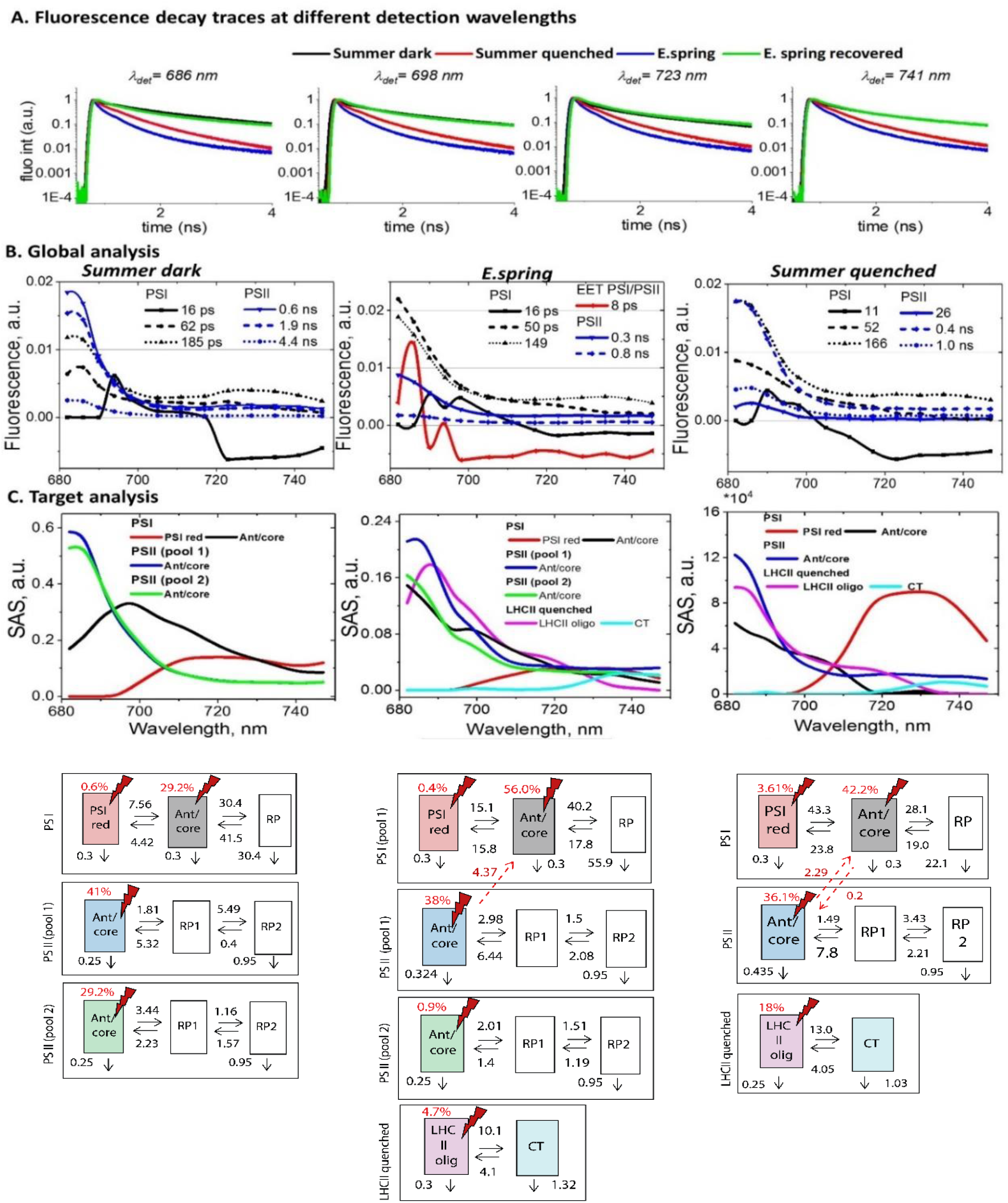
Time-resolved fluorescence of intact pine needles measured using TCSPC. (A) Fluorescence decay traces measured at −20 °C and at four characteristic wavelengths: 686 nm (mainly PSII, LHCII contributions), 698 nm (PSII, PSI contributions), 723 nm (mainly PSI contribution), and 741 nm (mainly PSI contribution).(B)Global analysis of pine needles in three states: Summer dark (S, dark-adapted summer needles, left row), ES (E.spring needles with “sustained quenching” present, middle row), Summer quenched (SQ, right row) (C) Kinetic target analysis of pine needles in the three states. The kinetic target analysis (SAS top, kinetic model with rate constants in ns^-1^, bottom) shows the results of the detailed target modeling of the fluorescence kinetics of pine needles. The rate constants (ns^-1^) were determined from global target analysis. Species-associated emission spectra (SAS) resulted from the fit of the target kinetic model in the corresponding state. *Note* that fluorescence decay measurements below 680 nm to detect the decreasing short-wavelength part of the spectra were not possible due to the extremely high scattering of the pine needles. This has no effect however on the ability to distinguish the various lifetime components kinetically and spectrally.

### Global analysis of the of the fluorescence kinetics of pine needles

To get the first hints on the mechanism(s) underlying sustained quenching in early spring global analysis^18^ was performed on all datasets: ES, S, SQ (Fig. 3 B) and ESR (SI 4.2) needles. Six Decay Associated Spectra (DAS) were required to fit the fluorescence kinetics in all four cases. Three DAS were tentatively assigned to PSI: 11-16 ps, 50-60 ps, 150-180 ps (Fig 3B, SI 4.1). Their spectra and lifetimes are reminiscent of previously reported PSI-related DAS in other plant species in vivo^16,17,19^. The fastest component represents energy equilibration between Chl_red_ and Chl _blue_^20,21^. The second DAS in S needles peaks around 685-690 nm with a broad emission in the 700-720 nm region. Therefore, it mainly represents PSI core decay in combination with the 710-720 Chls of Lhcas. The third DAS (185 ps) peaks at 680 nm with some contribution at 730 nm, and therefore, it should be primarily assigned to LHCII and some Chl_red_ in PSI. In quenched states (SQ and ES, in particular) these two DAS have substantially higher contribution around 680 nm and, as a result, the reconstructed steady-state PSI spectra have higher emission around 680-685 nm as compared to the unquenched states (S and ESR see SI 4.2). It indicates that upon quenching PSII and/or LHCII kinetics also contribute to these PSI-related components. In S and ESR states, the remaining three DAS peak at 682 nm and therefore, are assigned to PSII (Fig 3B and SI 4.1). Their lifetimes are 0.5-0.6 ns, 1.5-2.0 ns and 4 ns, similar to what was resolved previously for PSII in the closed state^16,17^. However, in ES two considerably shorter-lived PSII-related DAS were observed with lifetimes of 0.3 ns and 0.8 ns. This demonstrates that PSII kinetics is indeed quenched strongly in ES needles. Additionally, we resolved a DAS with 8 ps lifetime in the ES sample representing excitation energy transfer (EET) from Chl a pools emitting at ∼680 nm to those emitting at 700-730 nm, respectively. This indicates that there occurs efficient EET between PSII and/or LHCII and PSI in the ES needles.

In a nutshell, global analysis results show that sustained quenching in ES state involves both PSII/LHCII and PSI through a direct energy transfer. This mechanism provides much stronger quenching than the high-light-induced quenching in the summer needles (SQ).

### Target modeling of the fluorescence kinetics of unquenched pine needles

To resolve the mechanism(s) responsible for the ‘sustained quenching’ in pine needles in detail we performed target analysis^22^ of all the kinetic data. By testing kinetic models that had been validated previously on isolated subcomplexes^23–25^ it was possible to separate the PSI from the PSII kinetics. In S needles (Fig 4C, Summer dark) the PSI kinetics was described with two emitting compartments (PSI red, Ant/core) and one non-emitting radical pair (RP, Fig 3C). The resolved rates and Species Associated Spectra (SAS) are strongly reminiscent of the ones reported for the PSI-LHCI complex^25^. Kinetics from the PSII antenna/RC complex was described by one emitting compartment (Ant/core) and two non-emitting radical pairs (RP1 and RP2, Fig 3C) as reported previously^16,23,24,26^. According to their SAS and rates, the emitting species represents the equilibrated excited state of the PSII-LHCII super-complex. In our analysis, to achieve a very good fit, two pools of PSII complexes with different decay rates were required (pool 1, pool 2, Fig 3C, see the detailed comparison of the residuals plots in SI 4.3) for the S sample. The two pools differ in the rate constants of charge separation in a manner that is typical for PSII particles with different antenna sizes (see ^18,19^ for discussion and explanation), which result in different average lifetimes (Table 3 SM). In the S sample, PSI with its antenna accounts for 30% of the total absorption cross-section at the excitation wavelength (662 nm), while the two PSII pools account for 40% and 30% of the total absorption cross-section. No internal quenching related to any NPQ process of these PSII compartments was observed.

**Figure 4.**
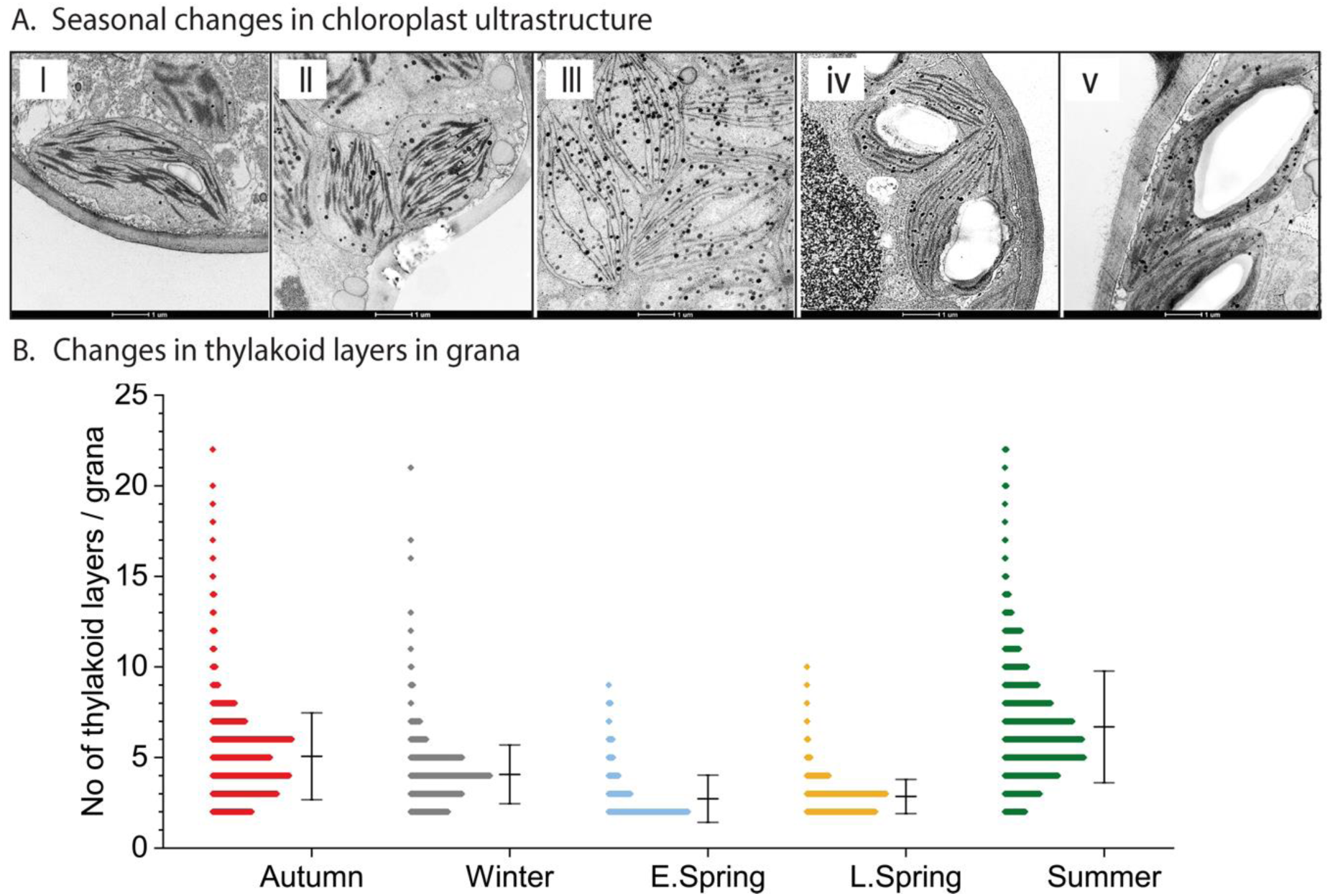
Transmission electron microphotographs depicting seasonal variations in chloroplast ultrastructure | in pine needles. **A**. Chloroplast structure in (I) Autumn (II) Winter (III) E. Spring (IV) L. Spring and (V) Summer. **B**. Histograms of frequency distributions of numbers of thylakoids per granum during the five distinct seasonal periods. The histograms were calculated from 80-100 electron micrographs per season, representing 2-3 chloroplasts per image (Error bars represent SD calculated from 800-1000 stacks per season).

The target model for the ESR needles is rather similar to that of S needles (SI 4.2). The main difference is that the smaller PSII pool (with a very small absorption cross-section of ca. 10%) shows an unusually small charge separation rate. The most likely explanation of this phenomenon is that a small percentage of PSII RCs has been photo inhibited during the 48 h relaxation process from the sustained quenching state, which agrees with NPQ measurements.

### Direct energetic transfer between PSII and PSI is the dominating component of “sustained quenching”

As for the S samples, also the description of the ES sample (Fig. 3C) required a three - compartmental target model for PSI and the presence of two PSII pools, each described by a three-compartmental model. The two PSII pools differ in the rate constants of charge separation in a similar manner as in S samples. The kinetics are also very similar to that of the S samples, however with a slightly increased charge separation rate. This can be interpreted as a consequence of partial reduction in the physical antenna size. This goes in line with an increase in the Chl a/b ratio in ES as compared to S samples (SI 6 Table 2).

The striking feature is the presence of a very strong contribution of a direct energy transfer from PSII to PSI. Without allowing for this direct energy transfer, the data could not at all be fitted adequately (SI 4.3, 4.4). This process has a transfer rate of 4.4 ns^-1^. A small compartment representing quenched and functionally detached LHCII was also needed for a satisfactory description of the ES samples (See more details on this component in the next section). Overall, the target analysis shows that direct energy transfer from PSII to PSI complexes provides the dominant quenching mechanism of PSII complexes in ES needles causing the pronounced “sustained quenching”.

### Comparison of sustained quenching and light-induced quenching

To model the fluorescence kinetics of SQ needles (Fig 4C) again three-compartmental models for PSI and PSII were required. However, unlike in previous cases, one single PSII pool was sufficient to describe the PSII kinetics.

Besides PSII and PSI, one additional component was needed to satisfactorily describe the fluorescence kinetics in the SQ samples. The rates, spectra and average lifetime (0.4 ns, Table 3 SI) of these compartments are strongly reminiscent of what was reported previously for highly quenched LHCII aggregates^27^, and the quenched LHCII complexes under NPQ conditions in wild type *Arabidopsis*^16^ and in *Arabidopsis* mutants depleted of the reaction centers^28^. We therefore assign this highly quenched component to the quenched LHCII antenna energetically disconnected from both PSII and PSI. However, the addition of this quenched LHCII compartment was not sufficient to fully describe the fluorescence kinetics (SI 4.4). The species-associated spectra (SAS) and kinetics of PSII still contained a substantial amount of 690-700 and 730 nm emission, entirely uncharacteristic for PSII super-complexes, but characteristic of PSI complexes^21,25^. Interestingly, also the SAS of the PSI antenna/core complex contains a 680 nm emission, which is characteristic for PSII. This strongly suggests – as can be directly deduced from the spectral properties - the presence of a direct energetic contact between PSI and PSII (Fig 3C). Introducing such a connection in the target model indeed resulted in a considerably better fit (χ^2^ decreased from 1.198 to 1.093, see SI 4.3). Also, the PSII and PSI kinetics became well-separated in such a model. The resolved direct energy transfer rate from PSII to PSI is quite high (2.3 ns^-1^), implying that a part of energy absorbed by PSII is actually funneled to PSI. However, this rate of direct energy transfer in SQ needles is only about half of that in ES needles, suggesting, that unlike in cold-induced sustained quenching, upon light illumination, spillover is not the major process. Contribution of the quenched antenna, on the other hand plays the major role.

### Efficiency of direct energy transfer quenching in different conditions

By evaluating the kinetic information provided in Fig. 3C one can get a quantitative estimation of the degrees of light use efficiencies (for photosynthesis) on the one hand, and for photoprotection/quenching on the other hand, present in both PSII and PSI in the different situations (c.f. SI 4.5, SI Table 4). In S needles 70% of the total absorbed energy drives PSII charge separation and only 30% PSI charge separation. In the SQ needles the percentage of energy flowing into the PSII reaction center drops to 7.1%, while it increases to 67% for PSI. Upon strong light illumination most of PSII reaction centers will be closed, and a fraction of that total energy flowing through the RC can produce harmful reactive species like ROS^29,30^. Since the percentage of energy flow into PSII decreases 10 times (70% in S and 7.1% in SQ) in SQ as compared to S, this means that the potential of oxidative damage is reduced by a factor of 10 by the quenching process. This factor is increased by a further factor of approx. 2 due to the large pool of functionally detached and quenched LHCII. So, in SQ needles the overall quenching provides a protection factor of 20 for PSII compared to S needles. In the sustained quenched ES samples this effect is much more extreme: Only 1.5% of the total absorbed energy in the system flows through the PSII reaction center potentially causing oxidative damage (assuming all other factors are equal). Thus, in total, ‘sustained quenching’ in ES samples provides a protection factor of approx. 40 to PSII (double of SQ) compared to S needles. This huge protection factor explains well why “sustained quenching” is so effective enabling pine needles to survive the harsh winter and early spring conditions.

### Massive destacking of grana membranes in the winter

In accordance with early reports^31,32^, we observed strong seasonal changes in chloroplast ultrastructure, including loss of grana stacks during early spring (Fig. 4A-III). Ultrastructural changes derived from morphometric analysis of electron micrographs (Fig. 4A) indicate that average numbers of grana per chloroplast steadily decreased from autumn to winter and reached minimal values in early spring, corresponding to just 68% of S values (SI 6 table 1). More strikingly, the number of appressed thylakoid layers/granum dramatically declined from A (4.97) to ES (2.72) and rose back in S (6.50) (Table 1). These changes corroborate the shift in grana stacking illustrated in Fig 4B. In S and A, 3-6 layered grana stacks accounted for 60-70% of total stacks. In contrast in ES samples, 70-75% and 15-20% of the grana had only two and three layers, respectively (Fig. 4B-II). These changes in ES samples were accompanied by a transient doubling of the number of lipid globules (plastoglobuli) per cell (SI 6, table 1) when the chloroplast structure deviated most strongly from the S state.

To obtain deeper understanding of the thylakoid plasticity in ES needles we also subjected needles collected at specific early spring and summer dates to EM analysis after artificially induced recovery under different light conditions (Fig. 5A-I to IV). When both S and ESR samples were exposed to HL for 30 (SQ1 for S samples, ESRQ1 for ESR samples) and 60 mins (SQ2 for S samples, ESRQ2 for ESR samples) similar destacking of grana (as of ES samples) have been observed (Fig 5B). These changes in grana structures recorded in ESR and S samples (Fig 5B-II) following short term HL exposure strongly corroborate the high dynamic plasticity of the thylakoid membrane, which appears to be essential for acclimation to the harsh boreal winters.

**Figure 5.**
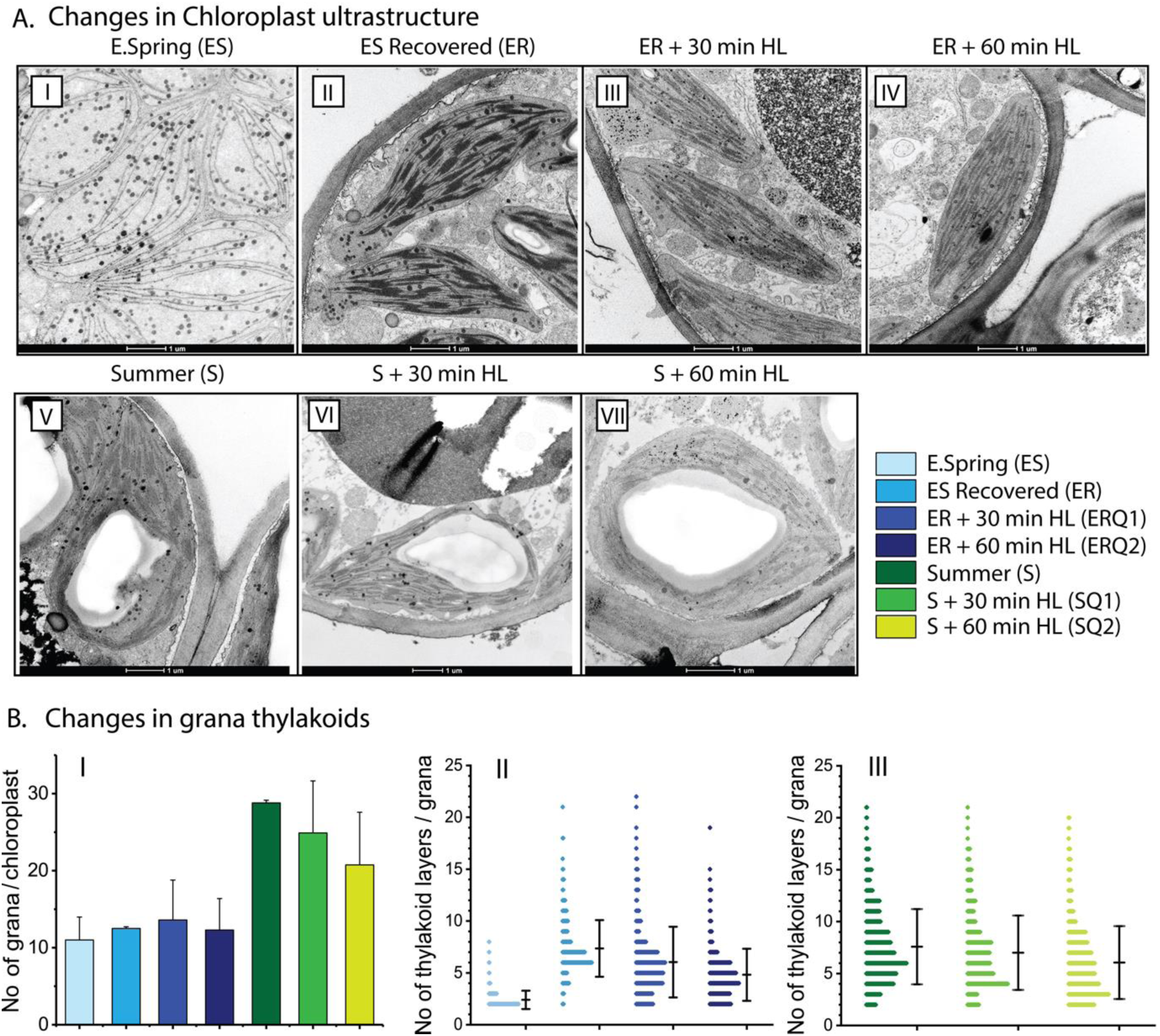
Artificial induction of changes in chloroplast ultrastructure of pine needles. **A**. Changes in chloroplast ultrastructure in (I) E. spring (ES), (II) E. spring samples recovered (ER) at 18oC for 48 hours with a photoperiod of 18 h at 80 μmol m^-2^s^-1^, (III) ER samples treated with 800 μmol m^-2^s^-1^ high light for 30 min (ERQ1), (IV) for 60 min (ERQ2). (V) Summer (S), (VI) Summer samples treated with 1200 μmol m^-2^s^-1^ high light for 30 min (SQ1), (IV) for 60 min (SQ2). **B**. The number of grana per chloroplasts (40 images typically containing 2-3 chloroplast per image) (I); Histograms of frequency distributions of numbers of thylakoids per granum in different (II) E. spring treated (III) Summer treated samples. Error bars represent SD obtained from analysis of 460-540 grana stacks.

### Changes in pigment and protein composition of ES, ER, SQ and S samples

We also analyzed pigments of the pine needles used for spectroscopic measurements and found substantial differences in pigment contents between S and ES needles (SI 6 table 2). In S and ES samples the Chl a/b ratios were 2.8 and 3.4, respectively, indicating that S needles had higher relative amounts of Lhcbs and other Chl b-binding proteins. The Chl /carotenoid ratio was also substantially lower in ES samples than in S samples, indicating that they had higher amounts of carotenoids (mainly lutein and violaxanthin/zeaxanthin). We observed no significant difference in total Chl pigment contents between S and SQ samples.

For quantifying the core and antenna proteins, isolated thylakoids were subjected to SDS-PAGE and were immunoblotted against specific antibodies (SI 5). This revealed that ES samples contain a lower abundance of core (PsaD, PsbD) and antenna (Lhcb2, Lhca4) proteins than in S, in line with earlier reports^33^ (SI 5). ESR samples contained lower amounts of PsbD and Lhcb2 compared to S, which might reflect on the minor amount of photoinhibition seen in fluorescence data.

## Discussion

Distinct changes in chloroplast ultrastructure, including loss of grana thylakoids and disorganization resulting in mostly single-/bi-stromal thylakoid membranes during the cold winter season, followed by recovery during the late spring/summer season have been observed in various conifers^8,13,32^. We observed similar strong seasonal variations in the chloroplast architecture of pine chloroplasts (Fig. 4, SI 6 table 1) and together with the fluorescence data presented above we are able to decipher the major molecular mechanism that connects these changes to the pronounced ‘sustained quenching’ described in conifers in the winter.

The seasonal reductions in maximal quantum efficiency of PSII (Fig. 1B-II) and light energy utilization during the winter (Fig. 1B-III) were associated with a substantial increase in excitation pressure, indicating overall reduction of the photosynthetic electron transport chain^34^. In early spring, conifers harvest light energy but use of the excitation energy for photosynthesis is severely restricted, resulting in potential photoinhibition. Hence, all overwintering conifers must develop highly efficient mechanisms for photoprotection of the photosynthetic apparatus. The major photoprotective mechanism for de-excitation of excess light energy in green plants and algae is the rapidly/ reversible ΔpH-dependent non-photochemical quenching (qE) that occurs in the pigment bed of LHCII proteins^3,16,35^. However, several alternative and/or supplementary photoprotective mechanisms for effective thermal deactivation of excess light energy have also been proposed^5,12,13,36–38^.

At first glance, the sharp decline in ΦPSII during early spring (Fig. 1B-III), might suggest that excess irradiance is dissipated via qE, which has been established as one of the most effective mechanisms for coping with the excess energy flux^3^. However, fast inducible qE was extremely low (Fig 1C-II,III) in ES samples and was not restored to S levels even after 48 h of recovery treatment. (Fig 1D-III). This is not unexpected, since, in contrast to the fast energy dissipation (qE) depending on ΔpH and PsbS-protein^3^ the energy dissipation during winter (sustained quenching) occurs under strong down-regulation, even perhaps in the absence of ΔpH^8,10,39^. However, we can assign this relatively small qE contribution to quenching to the small component (accounting for 4.7% of the absorption cross-section) of detached and quenched LHCII. This fraction was much smaller than in light-quenched samples (18%), and absent in both the dark-adapted S samples and recovered ES samples. Incidentally, a so-called ‘cold hard band’ has been reported in low temperature fluorescence spectra and kinetics of cold-acclimated evergreens^39,40^. Features of this band are reminiscent of quenched LHCII aggregates at low temperature^41^, hence it is probably correlated with the fraction of detached and quenched LHCII observed in ‘sustained quenching’ conditions. This led us to conclude that this quenching seen in conifers is not only ‘Sustained NPQ’ but also has a different mechanism.

Detailed quantitative energy partitioning analysis of total absorbed light energy by PSII demonstrates a strong increase (4 fold) of the fraction of constitutive thermal dissipation (ΦNO)^42,43^ (Fig 1C-III). Usually, this thermal dissipation (ΦNO) contribution is not explained by any clear mechanisms. However, in the case of ES samples it is strongly suggested by the data that the high levels of ΦNO do reflect the ΔpH-independent “sustained” energy quenching^8,10,39^. Our lifetime data provide an explanation for the origin of this component: A very high rate of direct energy transfer from PSII to PSI, provides this delta-pH independent (non-qE) quenching of PSII. Other phtoto protective mechanisms proposed, such as, role of photo-inactivated PSII complexes may also effectively dissipate excess excitation energy as heat^44^, and since the photoinhibitory quenching (qI), dependent on inactivation and/or degradation of D1, relaxes within hours (or longer), the process could also be considered a form of ‘sustained quenching’. Moreover, the involvement of RC quenching, based on enhanced S2QA- and S2QB-charge recombination, favoring non-radiative PSII RC dissipation of excess light, has been suggested to supplement or even replace the fast component of non-photochemical quenching during winter, and thus play a significant role in overwintering conifers^36,37^. However, we found no indication of photo inhibited PSII centers in the unrelaxed ES samples that displayed sustained quenching. A component with these characteristics was not observed in the ES samples. However, even if it would be present it could not explain the strong quenching effect since this component has longer average lifetime and higher fluorescence yield than the intact PSII units. In addition, we found no experimental evidence for PSII RC quenching either. All rate constants obtained from target modeling of the PSII antenna/RC pools in both unquenched and quenched states are quite normal. Thus, neither photoinhibition nor RC quenching alone could explain the strong ‘sustained quenching’. Moreover, changes in Y(I) suggest a lowered activity of PSI, even though PSI activity was less affected than PSII. In the absence of carbon fixation, one would expect higher acceptor side limitation, but contrastingly we found Y(NA) less affected, and donor side limitation [Y(ND)] was very high (Compared to S). This makes sense if PSII directly transfers energy to PSI, making PSI an energy sink for PSII by reducing linear electron flow, and in turn causing PSI donor side limitation (Oxidised P700) to increase.

The detailed structural and functional data presented here enable inference of a coherent quenching/photoprotection mechanism involving structural rearrangements of the thylakoids. The model explains Scots pine’s acclimation to the combination of harsh freezing temperatures and high solar radiation that occurs during early spring in northern boreal forests (Fig. 6). Acclimation to early spring conditions (Fig 6-II) results in massive loss of grana stacks (destacking) and formation of uniform membrane structures of stromal thylakoids. This allows for a redistribution and randomization of PSII and PSI. The reduction in the spatial distance between PSII and PSI complexes increases the probability for direct energy transfer from PSII to PSI^45–47^. This protective mechanism severely restricts linear electron flow since it strongly quenches PSII activity. It also has strong similarities to the potent quenching observed in other extreme environmental conditions when PSII requires strong protection. Examples include drying-induced quenching in lichen^48^ and heat-induced quenching in *Symbiodinium* cells of coral^49^. In both cases, destacking and reorganization of the thylakoid membrane occur, resulting in proximity of PSII to PSI complexes with direct energy transfer.

**Figure 6.**
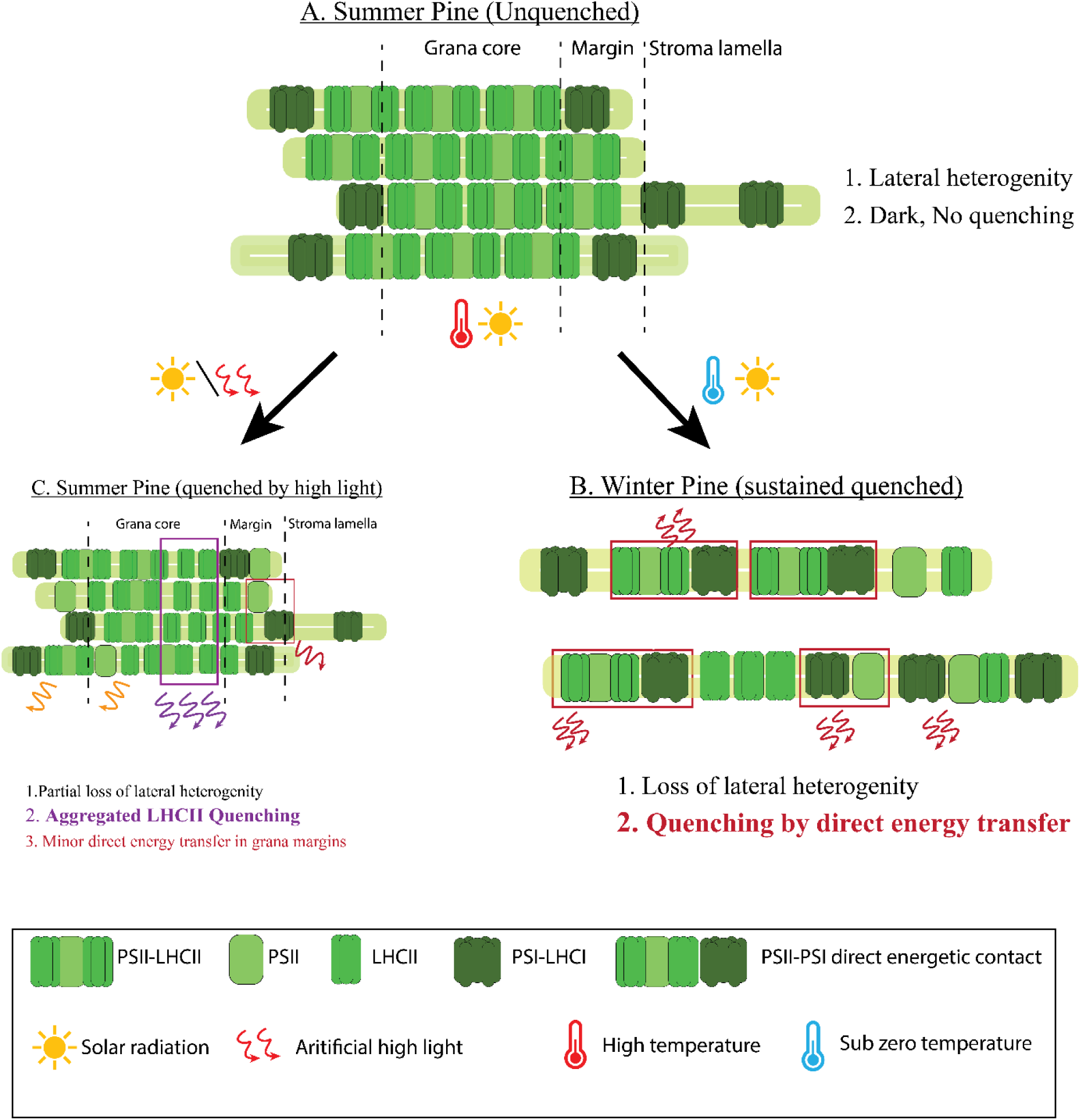
**Molecular model for acclimation of photosynthetic machinery under changing natural environmental conditions,** such as, in Summer unquenched (A); Winter quenched (B); Summer quenched (C) in Scots pine.

In conclusion, during early spring only a very small fraction of the absorbed light energy in PSII antennae is utilized for PSII photochemistry (ΦPSII) and the fast inducible NPQ (qE) is low since it not only requires a high ΔpH^3,6,16^ but there is a strong static quenching present already. On the other hand, thylakoid rearrangement and movement of complexes are likely coupled to the high zeaxanthin content^50^ (SI 6 Table 2), and probably to pronounced phosphorylation of antenna and core proteins present under these conditions^51^. In this situation, the excess energy is dissipated primarily through direct energy transfer from PSII to PSI, which appears in steady-state (PAM) fluorescence measurements as a non-regulated constitutive energy quenching. An additional, albeit smaller, amount of quenching and photoprotection is provided under sustained quenching conditions by the qE mechanism in functionally detached LHCII. It is possible that this fraction is linked to large aggregates containing LHCII, PsbS and other small proteins, previously detected in cold-adapted pine needles^52^. Thus, conifers appear to have a powerful protective mechanism that resembles those found in other photosynthetic organisms of relatively harsh environments^48,49^. This protective mechanism plays a crucial role in the survival of pine from harsh boreal winters.

## Material and methods

### Weather

Weather data were recorded from October 2015 to May 2016, from November 2016 to June 2017, and from October 2017 to June 2018 on an hourly basis at the weather station of the Sweden’s meteorological and hydrological institute, Umeå stations. (available on https://www.smhi.se).

### Plant material and sampling

Fully developed needles were collected frequently, from Oct 2015 to May 2016, from Nov 2016 to June 2017 and Oct 2017 to June 2018, inclusive, at midday (8.00-10.00) from fully exposed south-facing branches of five Scots pine (*Pinus sylvestris* L.) trees growing on Umeå University campus (63° 49’ N, 20° 18’ E). The collected needles were immediately stored in darkness, at 0-5°C, in a laboratory at Umeå Plant Science Centre, and subjected to measurements (described in the following sections) shortly after sampling to minimize any further changes in the photosynthetic machinery. For recovery treatment samples from winter-stressed branches were collected in mid-March and subjected to recovery treatment, involving exposure to very low light (80 μmol photons m^-2^s^-1^) at 20°C^52^. Further details are provided (together with other details regarding the sampling dates and procedures) in SI 1.

### Modulated chlorophyll fluorescence and (P700+) measurements

Chlorophyll fluorescence was measured using a Dual-PAM-100 instrument (Heinz Walz GmbH, Effeltrich, Germany) at room temperature (20°C) after 30 min dark adaptation at low temperature [below 5°C, winter samples (samples collected during November to May)] or room temperature [(20°C, summer samples (samples collected during June to October)]. Maximum photochemical efficiency of PSII was calculated as Fv/Fm = (Fm - Fo)/Fm. To perform all measurements, we chose to record a light response curve where we measured Fv/Fm first, followed by Pm (maximal P700 oxidation), and then changing the actinic light intensity stepwise from 33μE to 1469μE. During recording light response curve, samples were kept in one light intensity for 3 mins followed by a saturating pulse of 4000μE to get the optimal values. Partitioning of absorbed light energy was estimated as ΦPSII + ΦNPQ + ΦNO = 1^42,43^. The effective photochemical quantum yield of PSI, Y(I), was calculated as Y(I)=1-(Y(ND)+Y(NA))^53^. Here, Y(I) is the effective quantum yield of PSI (ΦPSI) when reaction centers (RCs) are open with reduced donor side (P700) and oxidized acceptor side (A); Y(NA) is the energy dissipation due to acceptor side limitation, reflecting the redox state of the acceptor side when RCs are closed with P700 reduced and both acceptors reduced; and Y(ND) is the energy dissipation due to donor side limitation, reflecting the redox state with both P700 and A oxidized.

### Ultrafast fluorescence measurements

Intact needles were subjected to ultrafast time-correlated single photon counting measurements at −20 °C in a rotation cuvette, as previously described^16,17^. Pine needles were immersed in 60% PEG solution (pH 7.5), which was used as an anti-freeze. Tests confirmed that this solution did not significantly affect either the Fv/Fm values or fluorescence kinetics. These conditions allowed maintenance of the physiological and quenching states for a long time during the measurements and prevention of radiation damage or further quenching induction by the very weak measuring light. Needles sampled in summer (S needles) were measured in both unquenched and fully light-quenched states. For measurement in the unquenched state, they were dark-adapted for 30 min, then quickly frozen to −20°C in the rotation cuvette. For measurement in the light-quenched state, the needles were illuminated by red light (intensity 1000 μmol m^-2^sec^-1^) for 30 min. at room temperature. During this time, fluorescence intensity was monitored and after 30 min. a steady-state level was reached. The needles were then frozen quickly under illumination in the rotation cuvette to - 20°C. Early spring samples (ES samples) were measured either in the original ‘sustained quenching state’ by transferring them directly at −20 °C into the cooled rotation cuvette (SI 4.1), or in a ‘recovery state’ induced by exposing them to very low light (80 μmol m^-2^s^-1^) at 20°C, as described earlier, then freezing them (as in the measurements of the summer needles in unquenched state). After each fluorescence lifetime measurement, the samples were fast-frozen in liquid nitrogen and stored at −80 °C for further pigment analysis. The laser settings for the measurements were 40-60 μW power, 4 MHz repetition rate, and 1 mm diameter beam size.

### Pigment and protein analysis

Pigments were extracted from frozen pine needles with 80% acetone. Chl a/b and Chl/carotenoid ratios were calculated by fitting the extracts’ absorption spectra with the spectra of the individual pigments, and the relative amounts of carotenoids were determined by HPLC as previously described^54^. For protein composition analysis intact thylakoids were isolated followed by Grebe et al. 2018^55^, subjected to SDS PAGE separation followed by Damkjær et al. 2009^56^ and immunoblotted against Anti PsbD, PsaD, Lhcb2 and Lhcb4 as per manufacturer’s instructions (Agrisera AB, Sweden).

### Transmission electron microscopy

Thin slices (< 0.5 mm) from the middle region of pine needles were cut in tap water and fixed in 4% paraformaldehyde, 2.5% glutaraldehyde (TAAB Laboratories, Aldermaston, England) in 0.1M or 0.05M (May and June) sodium cacodylate buffer, pH 7.4 (TAAB Laboratories, Aldermaston, England). Thoroughly washed samples were post-fixed in 1% osmium tetroxide (TAAB Laboratories, Aldermaston, England). The fixed material was dehydrated in ethanol series with increasing concentrations and propylene oxide and finally embedded in Spurr resin (TAAB Laboratories, Aldermaston, England). Ultrathin sections (70 nm) were post contrasted in uranyl acetate and Reynolds lead citrate and further examined with Talos 120C electron microscope (FEI, Eindhoven, The Netherlands) operating at 120kV. Micrographs were acquired with a Ceta 16M CCD camera (FEI, Eindhoven, The Netherlands) using TEM Image & Analysis software ver. 4.14 (FEI, Eindhoven, The Netherlands). The chloroplast ultrastructure was analyzed from the electron micrographs by measuring the average number of chloroplasts per cell, average number of grana per chloroplasts and average number of appressed thylakoids per grana stack (Ng)^57,58^.

### Statistical analysis

The significance of between-mean differences was assessed by t-tests and p values were recorded. One (*), two (**) and three (***) asterisks in the presented figures indicate P≤95%, P≤99%, and P≤99.90, respectively, for fluorescent and electron microscopic measurements.

## Acknowledgements

This work was supported by SE2B Horizon 2020 under grant agreement no. 675006 (SE2B) to SJ (Umeå University) and RC (Vrije Universiteit Amsterdam), the Swedish Research council (VR) and the Kempe foundation to SJ and the Netherlands Organization for Scientific Research (NWO-Vici grant to RC).

## Author Contributions

SJ and ARH conceived the idea; PB, AGI, ARH, and SJ designed the experiments; PB, ZZ, SP and TS performed the chlorophyll fluorescence and P700 experiments; PB performed TEM studies and protein quantification; PB and AGI analysed chlorophyll fluorescence data; VC performed ultrafast time-resolved fluorescence measurements; VC and ARH analyzed the time-resolved fluorescence data; PB, VC, AGI, ARH, RC and SJ analysed and discussed all results; PB, AGI, VC, ARH and SJ wrote the paper.

## Supplementary information

### Supplementary information 1. Sampling for seasonal profiling

**1A.**
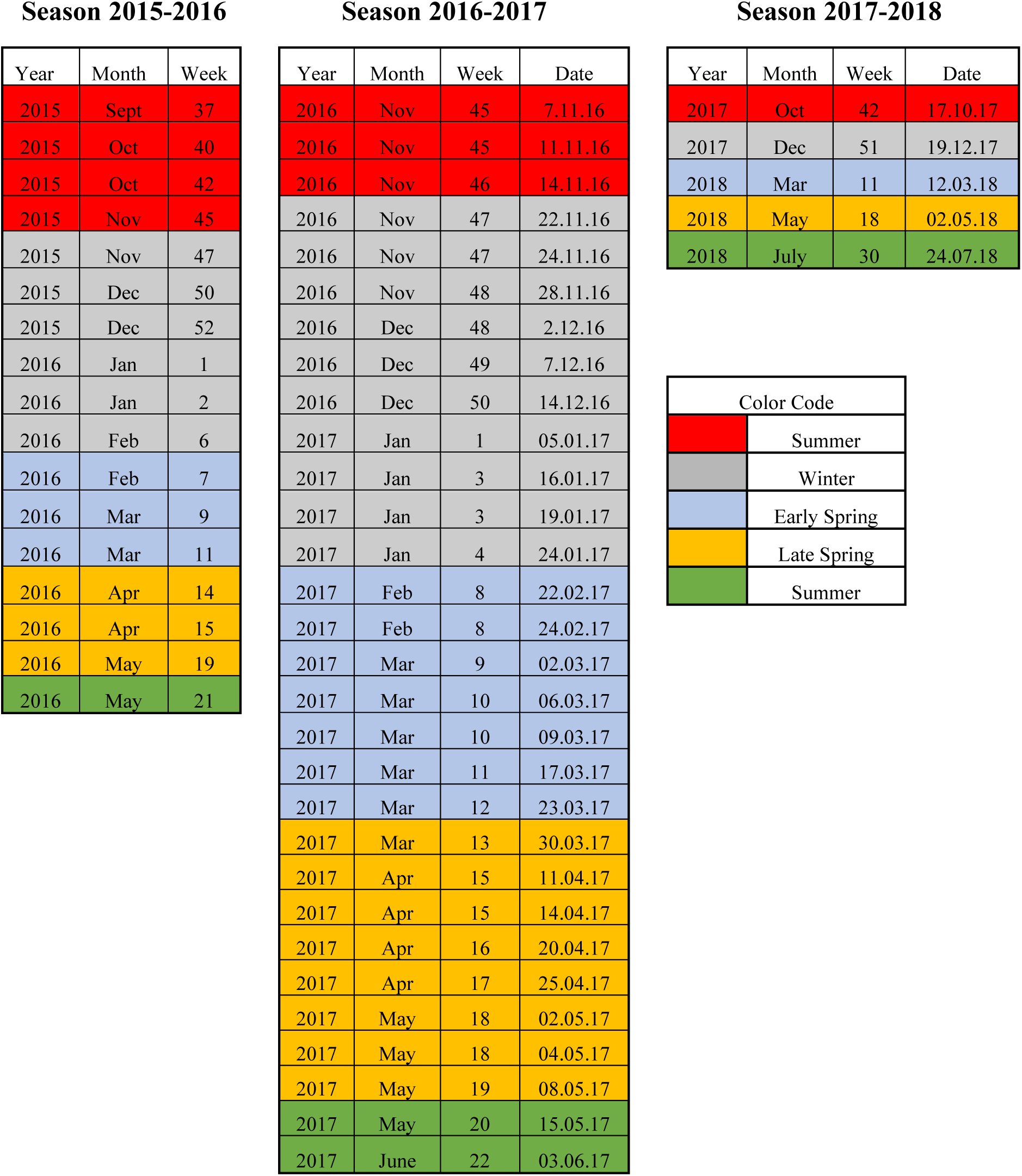
Sampling dates for Fluorescence (2015-2016, 2016-2017 and 2017-2018) and P700 measurements (only 2016-2017 and 2017-2018)

**1B.**
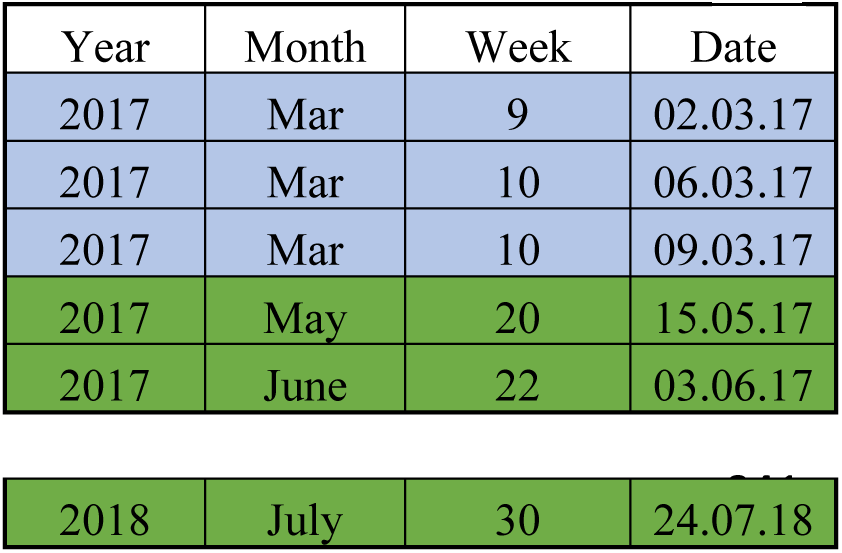
Sampling for Time resolved measurements (2016-2017 and 2017-2018)

**1C.**
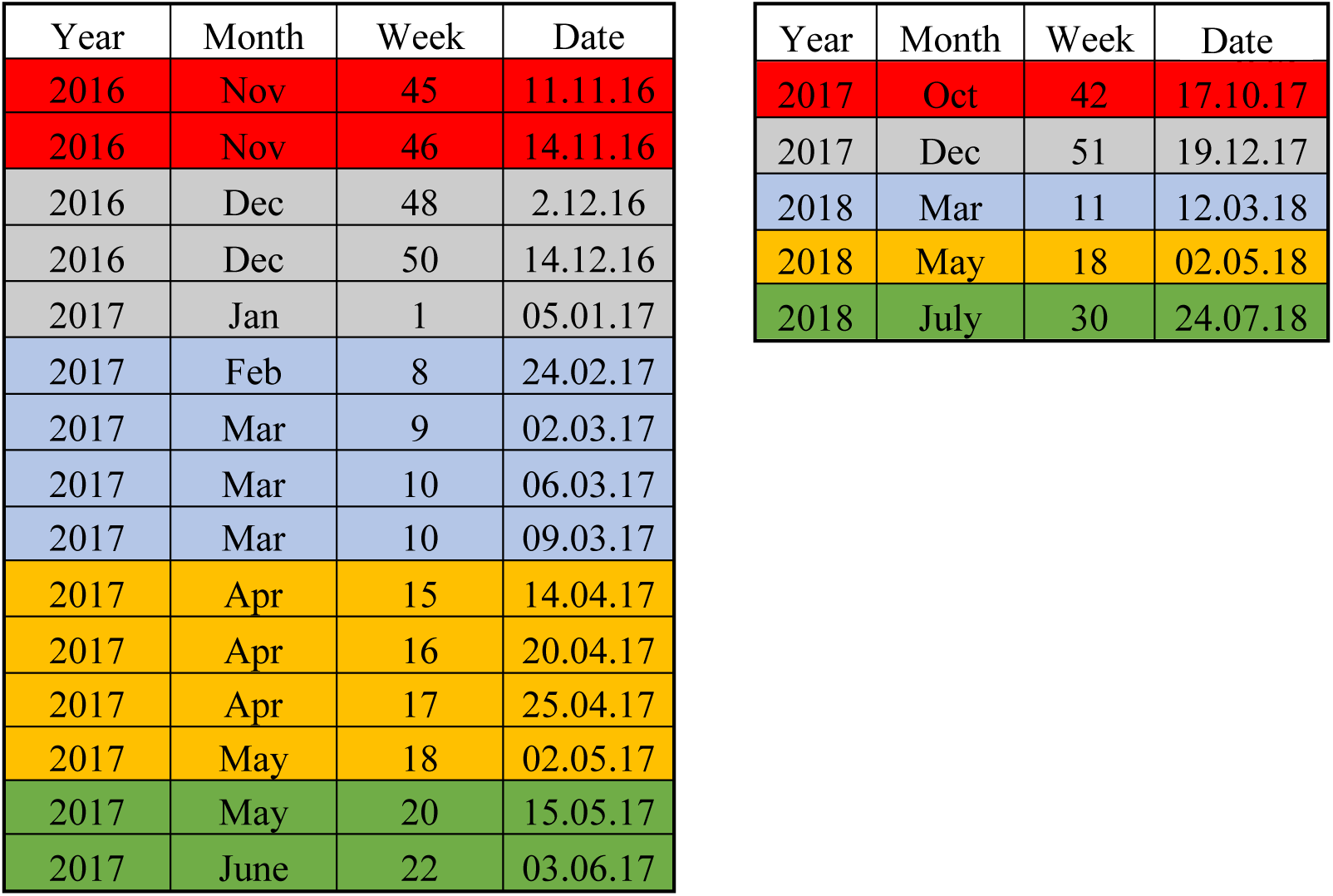
Sampling for seasonal Electron Microscopy (2016-2017 and 2017-2018)

**1D.**
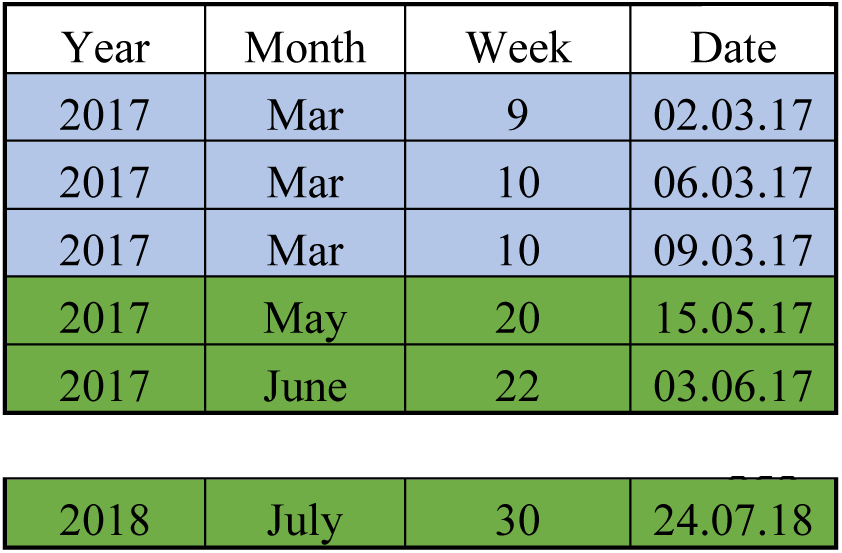
Sampling for protein quantification (2016-2017 and 2017-2018)

### Supplementary information 2. Seasonal performance of PSII

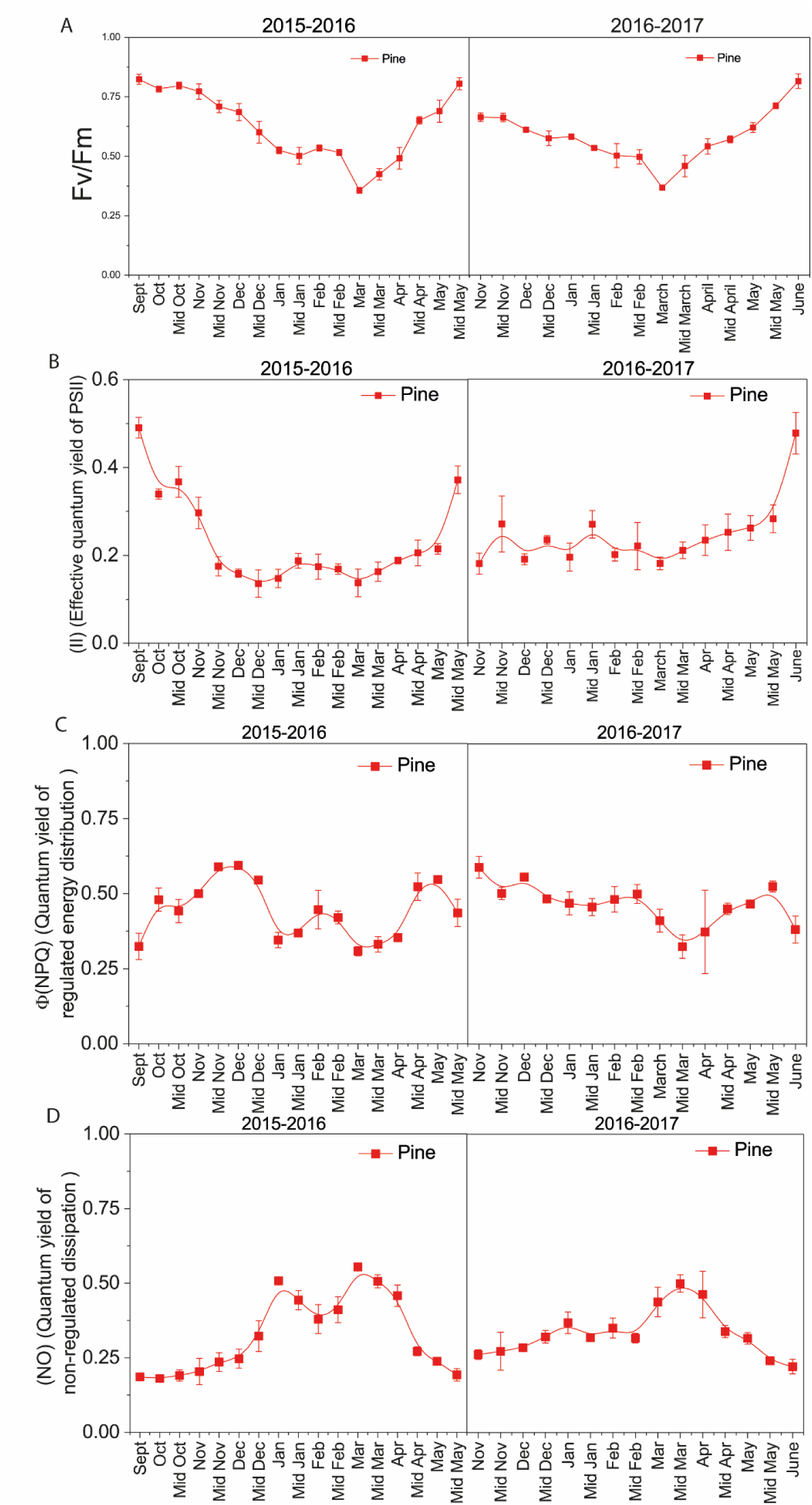

### Supplementary information 3. Seasonal performance of PSI

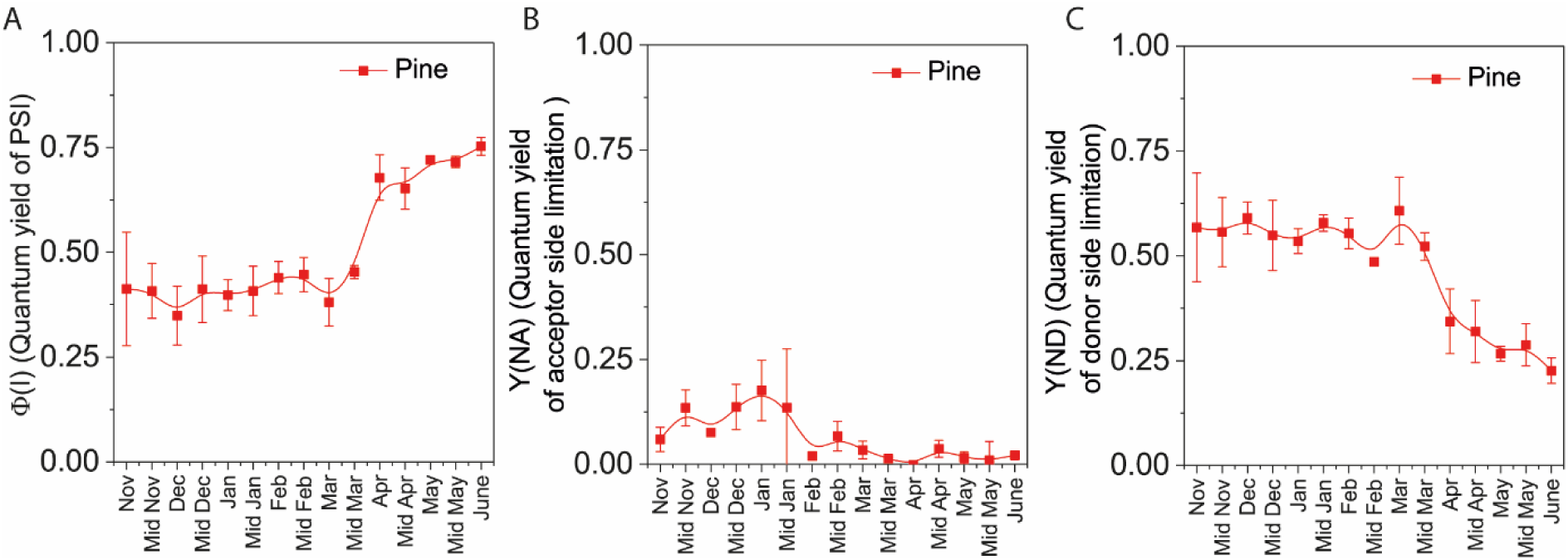

### Supplementary information 4. Lifetime measurements of pine needles

**4.1.**
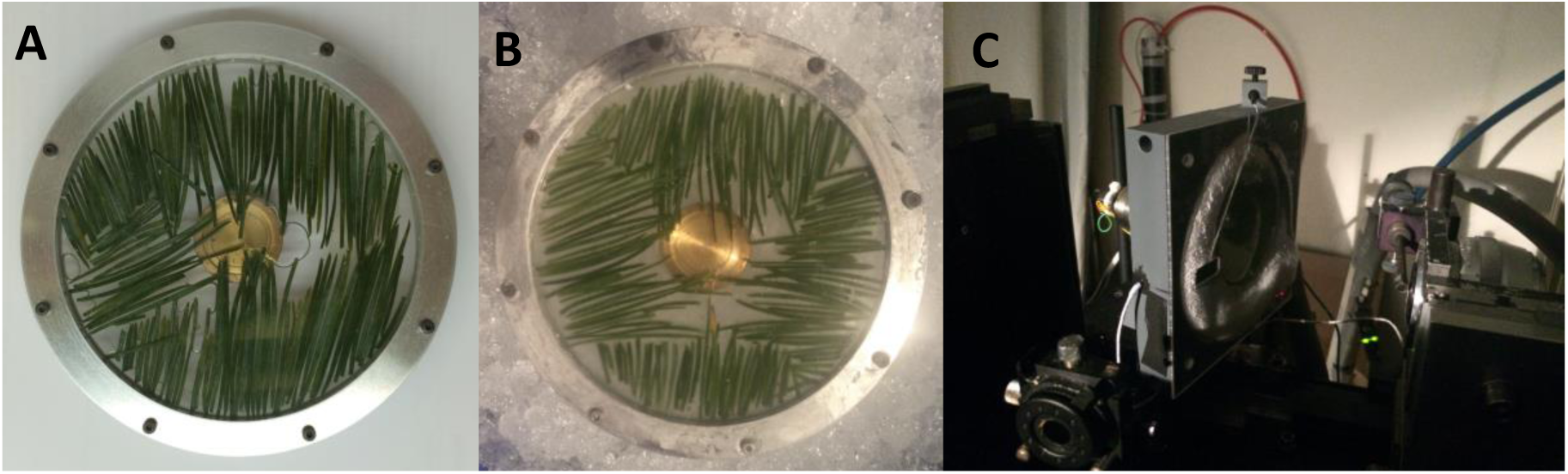
Measuring cuvette with pine needles inside in S state (A) or ES (B). Temperature control chamber, with the cuvette inside it during the experiment.

**4.2.**
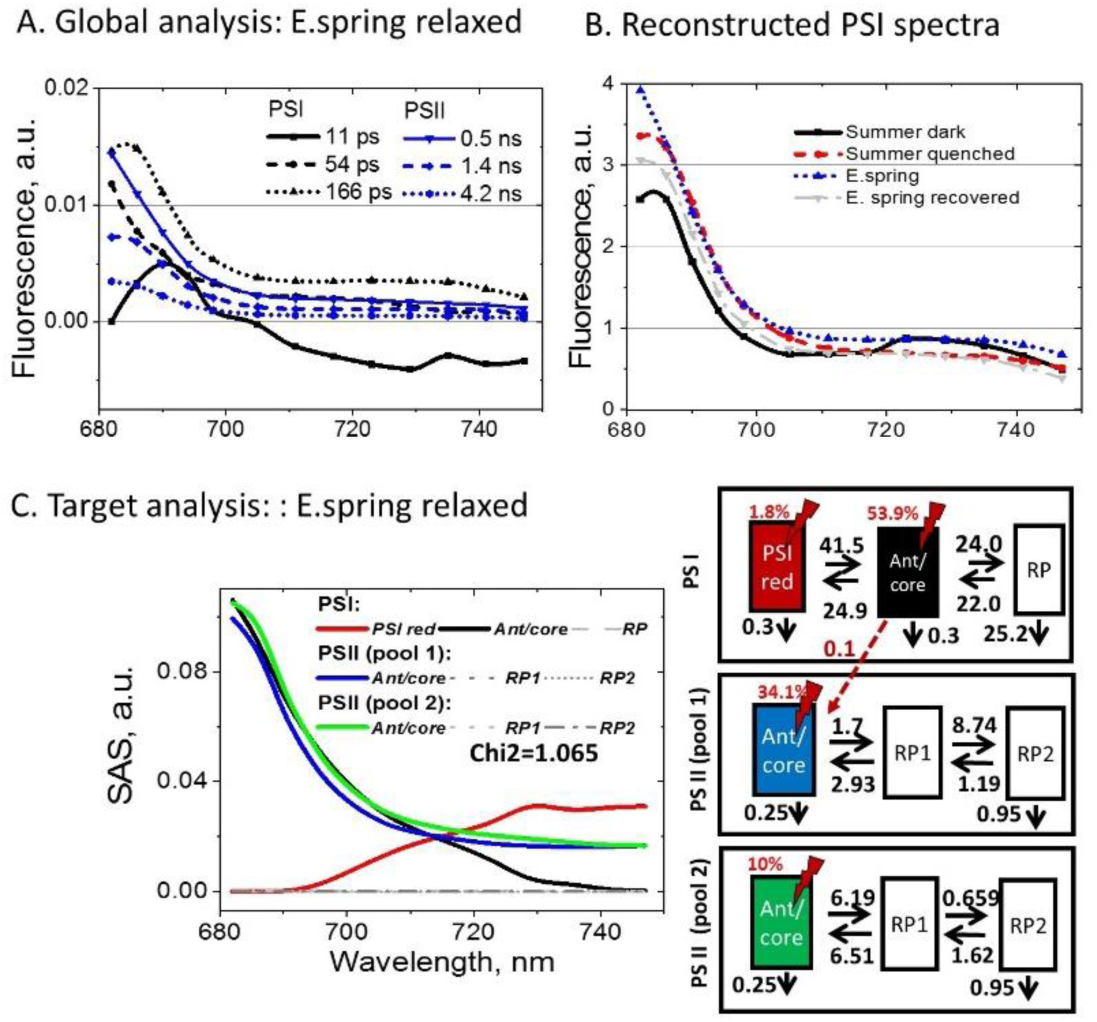
(A) and target (C) analysis of E.spring needles relaxed. (B) Reconstructed steady-state PSI spectra in four measured states

**4.3.**
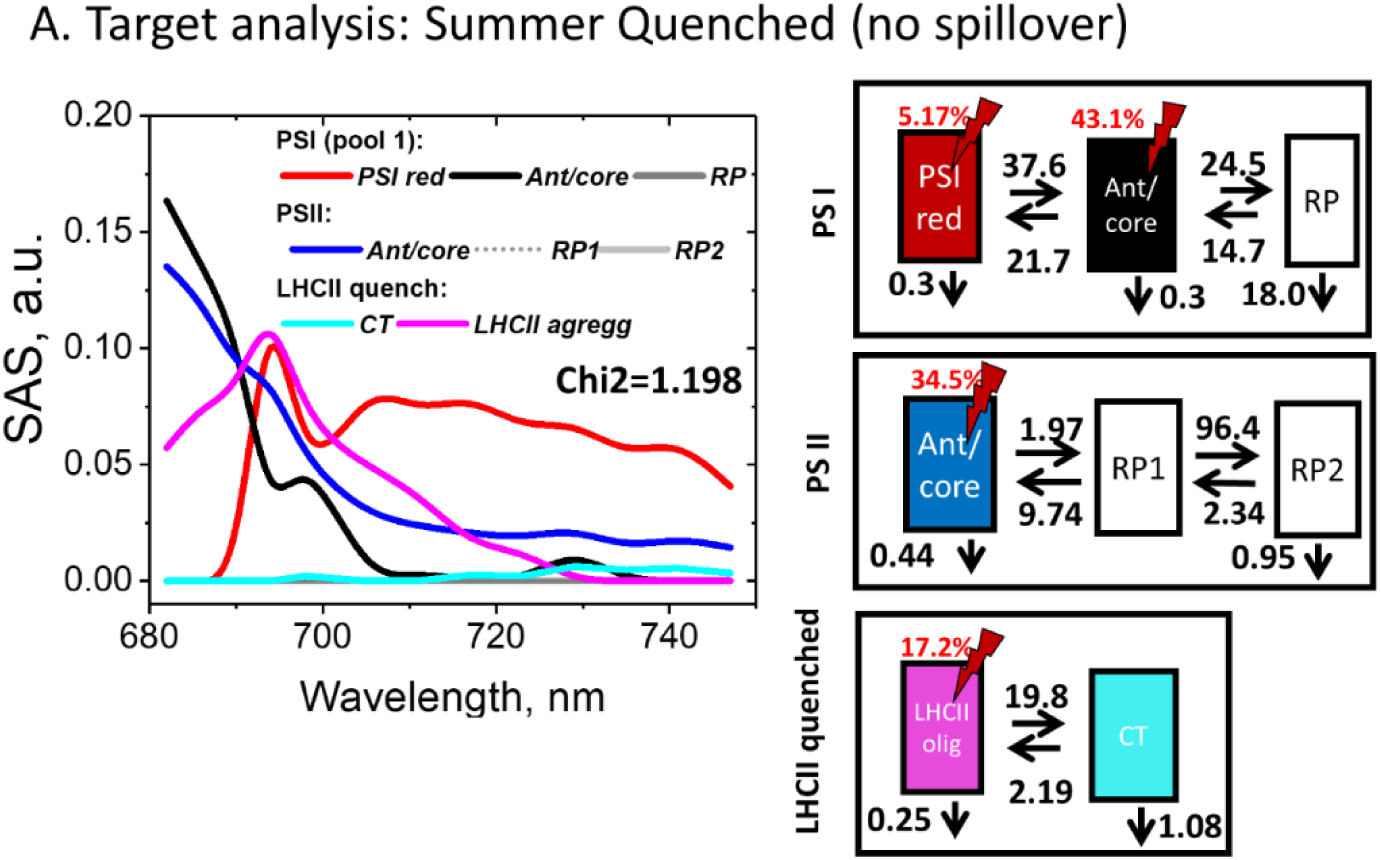
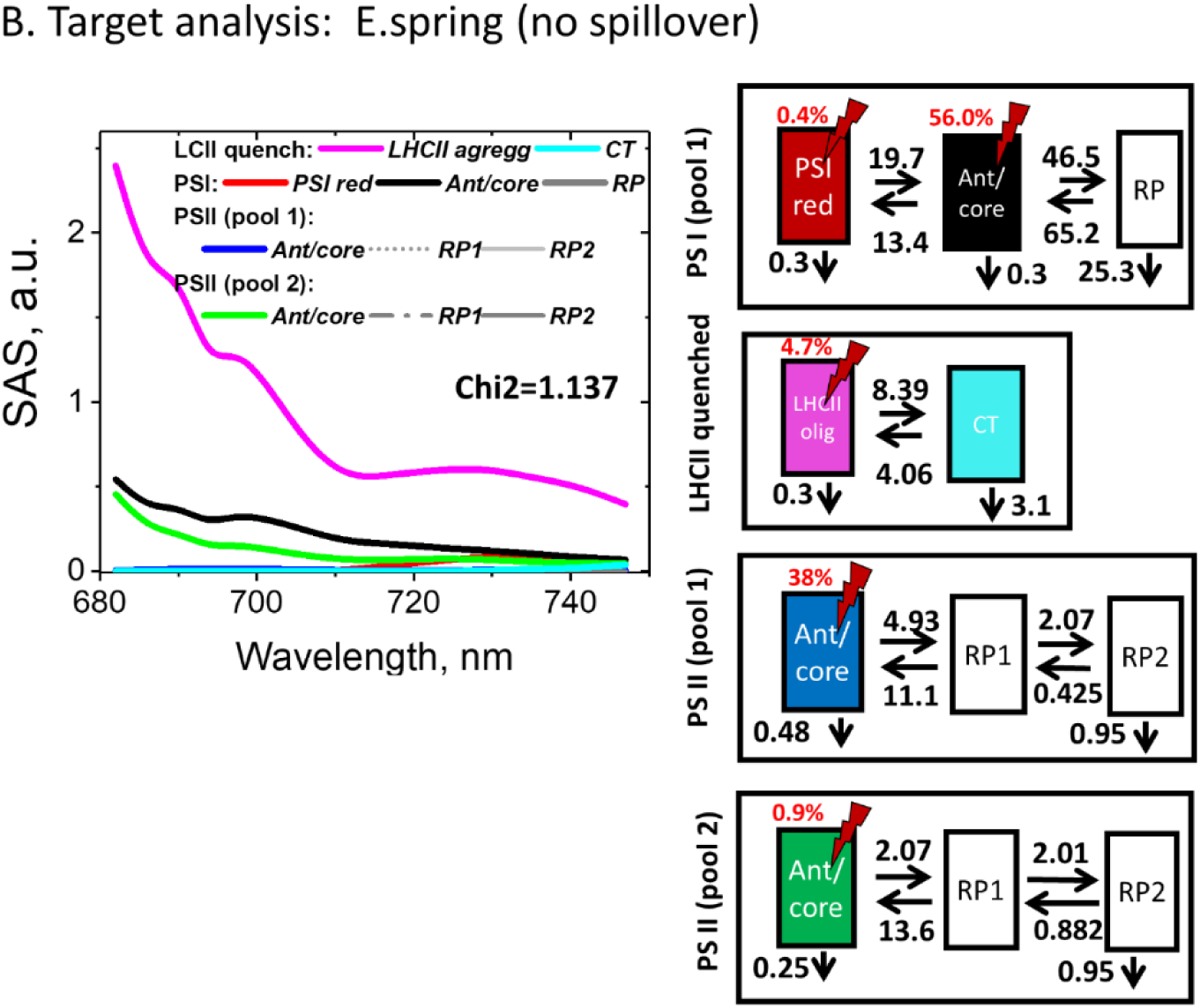
Targeted analysis of fluorescence kinetics of pine needles without spillover mechanism present

**4.4.**
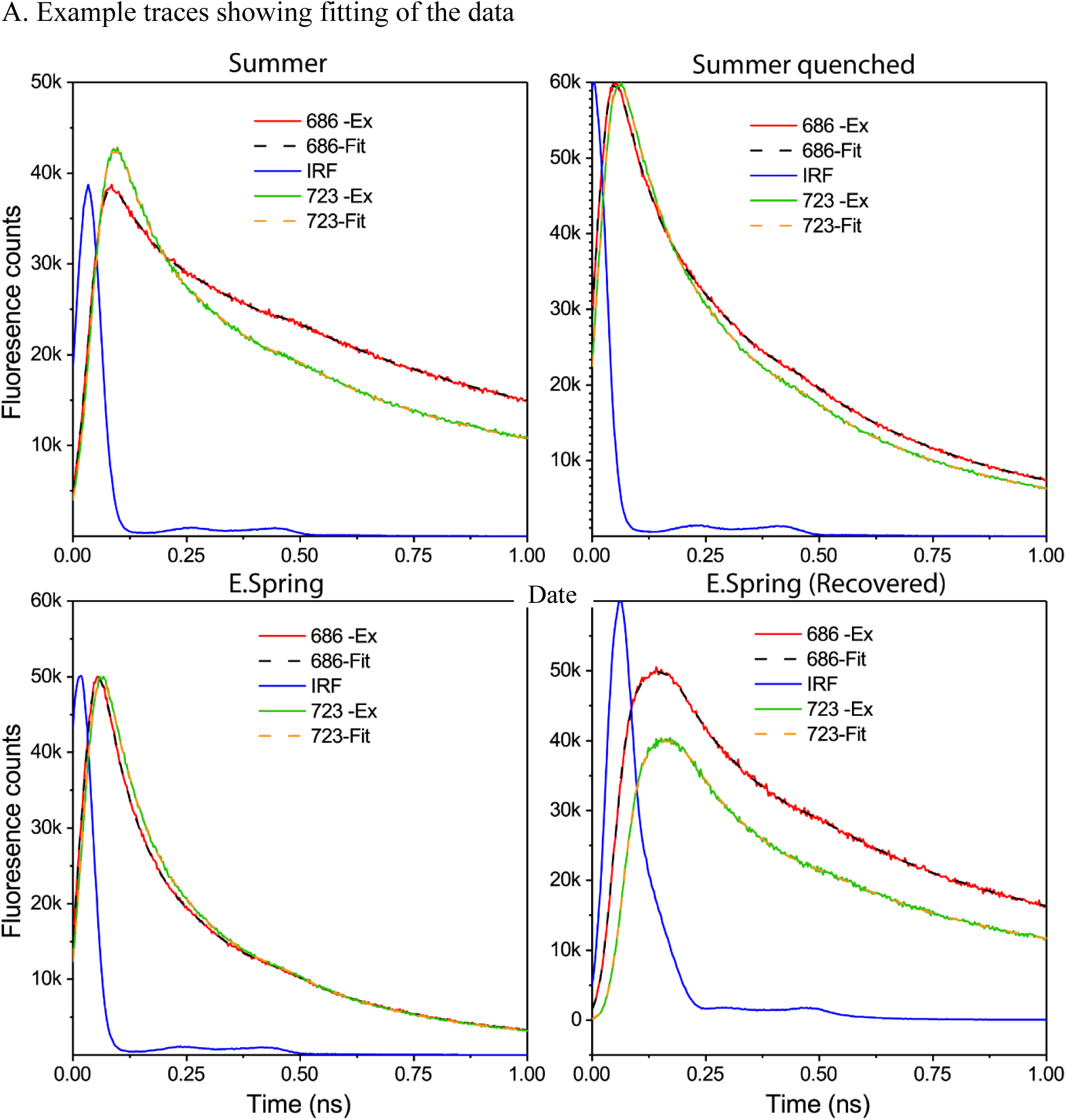
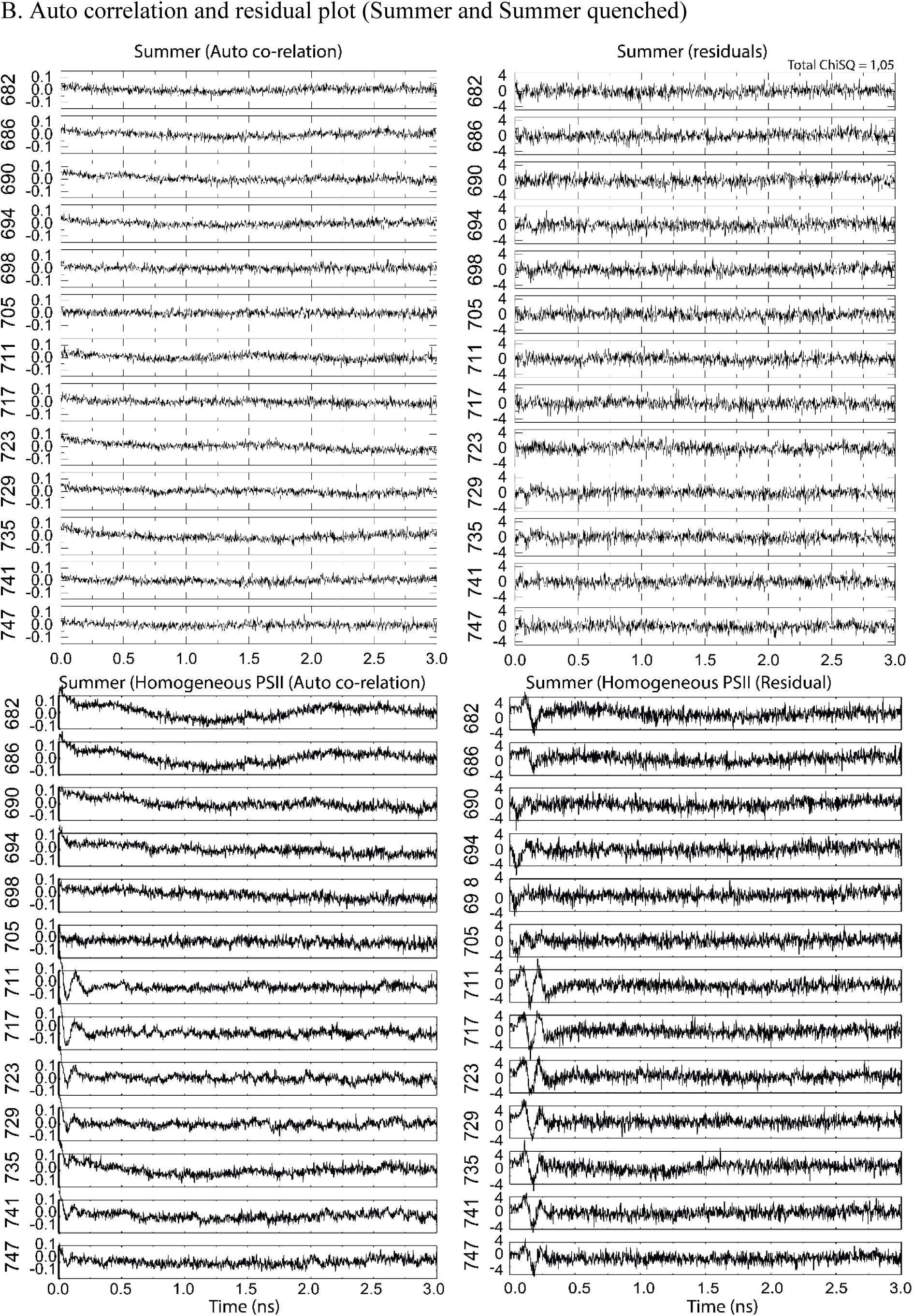
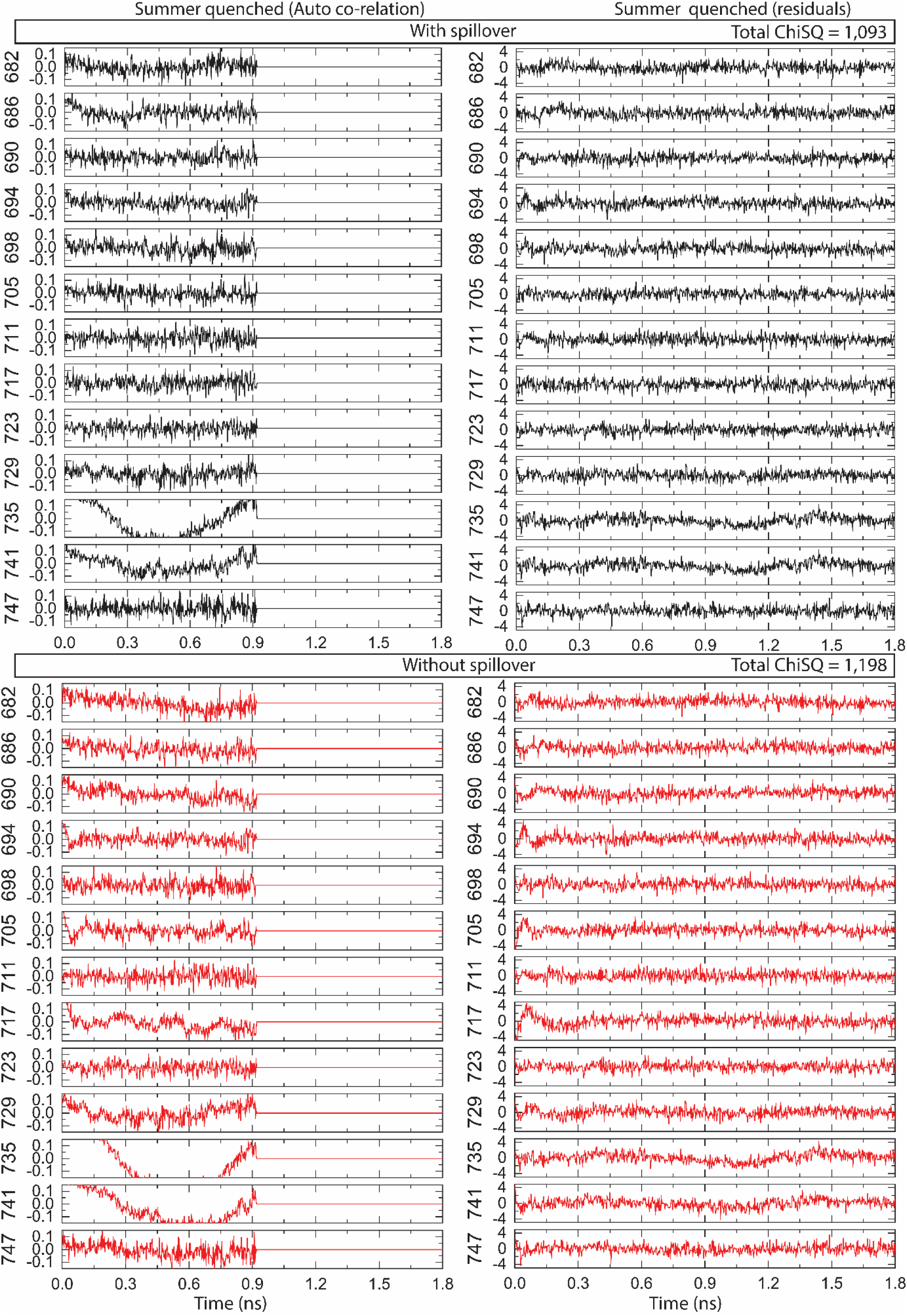
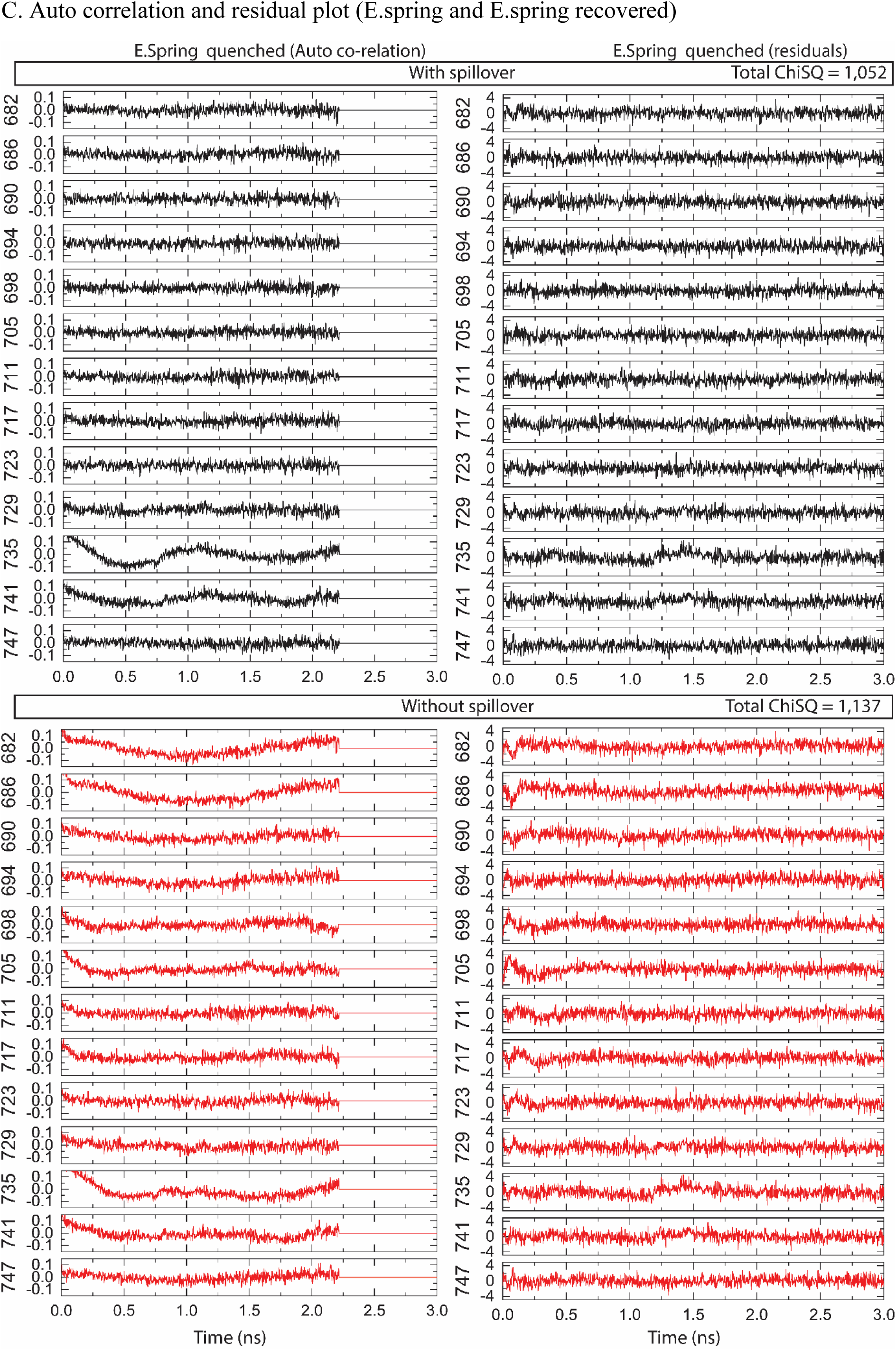
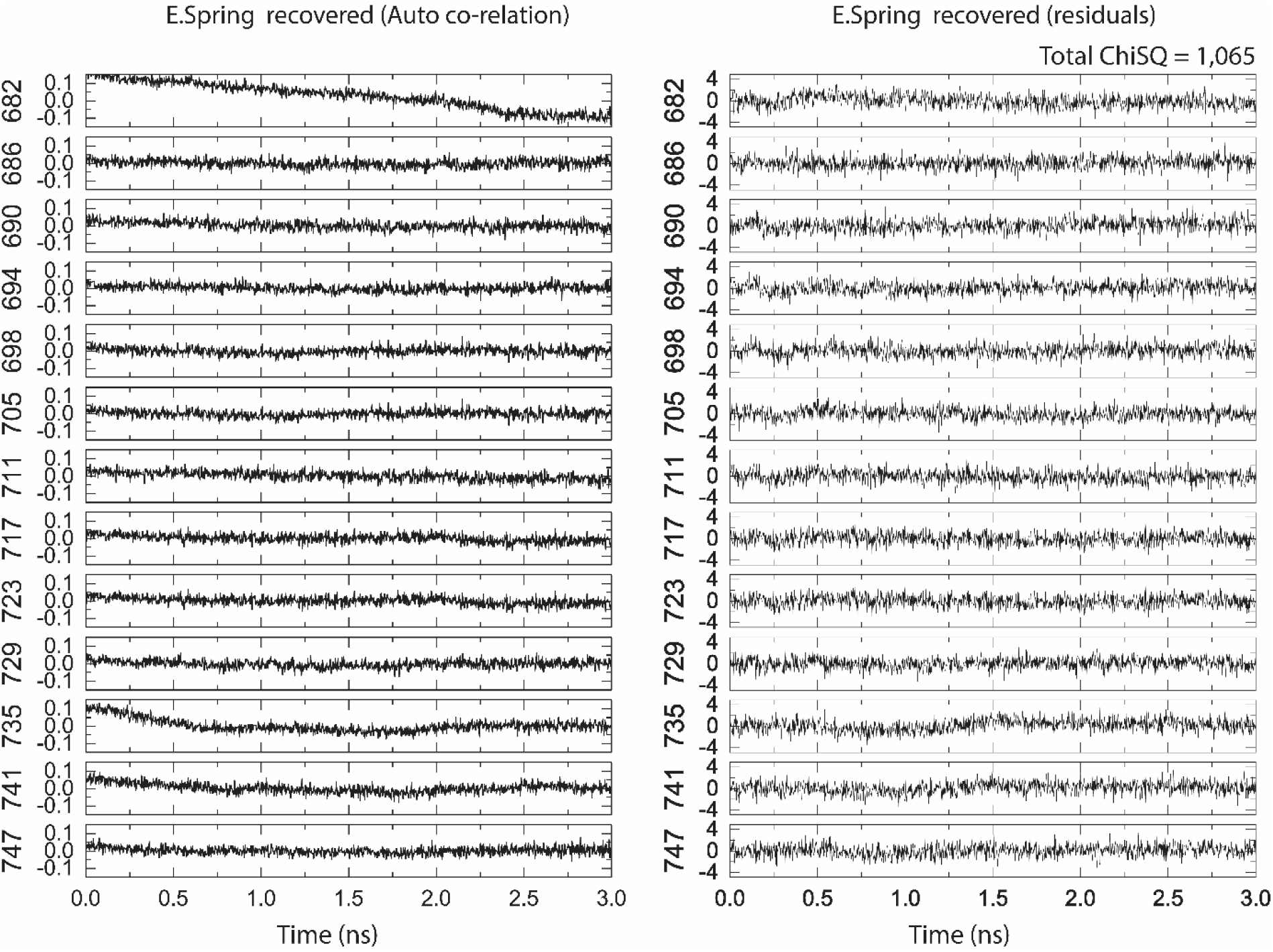
Auto co-relation and residuals plots A. Example traces showing fitting of the data B. Auto correlation and residual plot (Summer and Summer quenched) C. Auto correlation and residual plot (E.spring and E.spring recovered)

**4.5.**
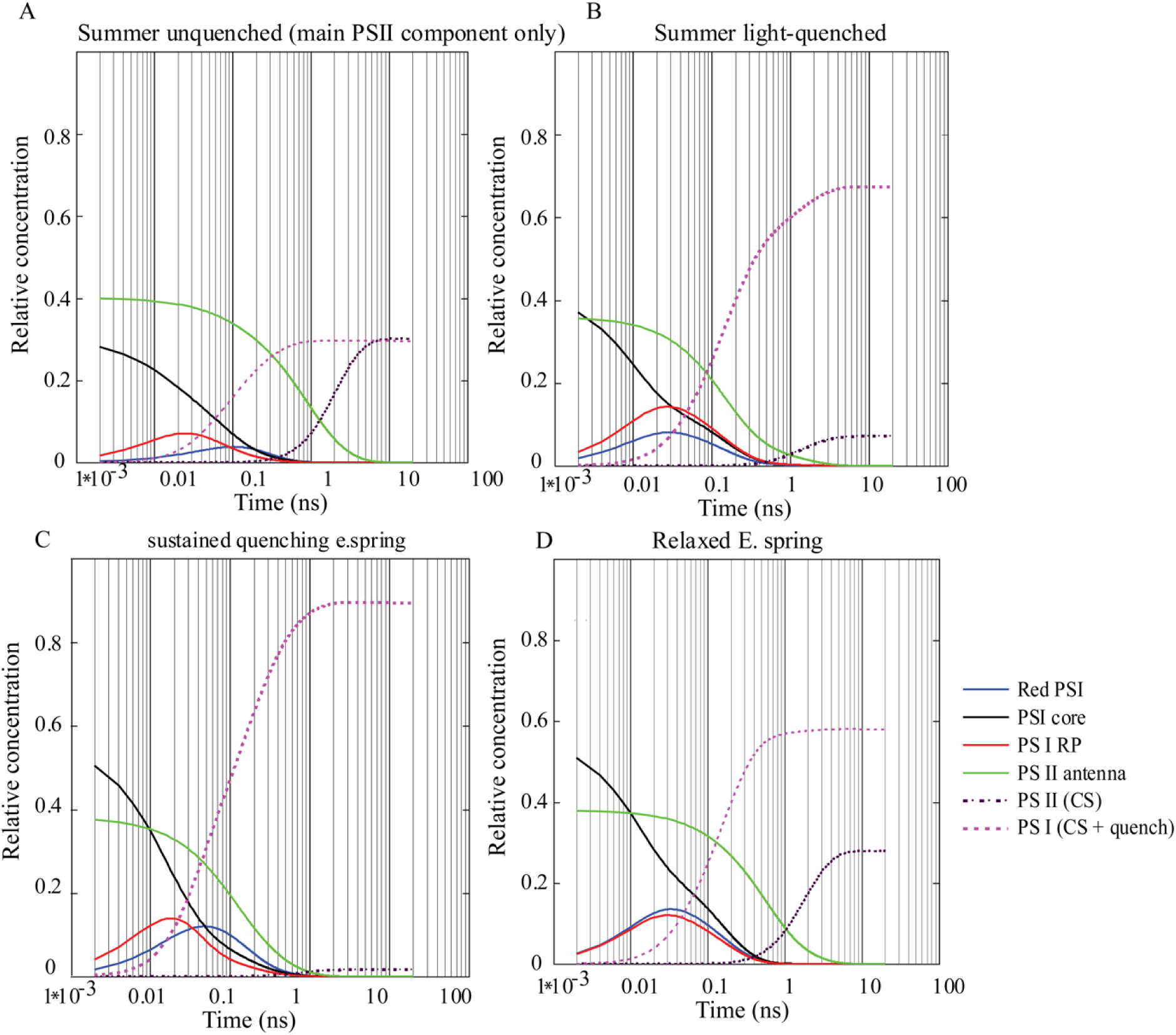
Time-dependent (on log time scale) populations of selected PSII and PSI compartments as calculated from the fluorescence kinetics (Fig. 3C). The dashed/dotted curves show the kinetics energy (normalized to thr total ablsprtion cross-section) flowing into PSI (purple dashed curves) and PSII (dotted black curves). The inital exitation input was taken from the excitation vectors of corresponding target analysis results (Fig 3C). Depending on the state of the respective reaction center, that energy will be either used for photochemistry or will be deactivated non-radiatively (quenching). See Table 4 SI for the percentages. Black (PSI) and green (PSII) curves show the time course of the excited state populations.

### Supplementary Information 5. I. SDS_PAGE separation of thylakoid proteins loaded based on equal chlorophyll. II. Quantification protein by specific antibodies against PsbD, Lhcb2, PsaD and Lhca4.

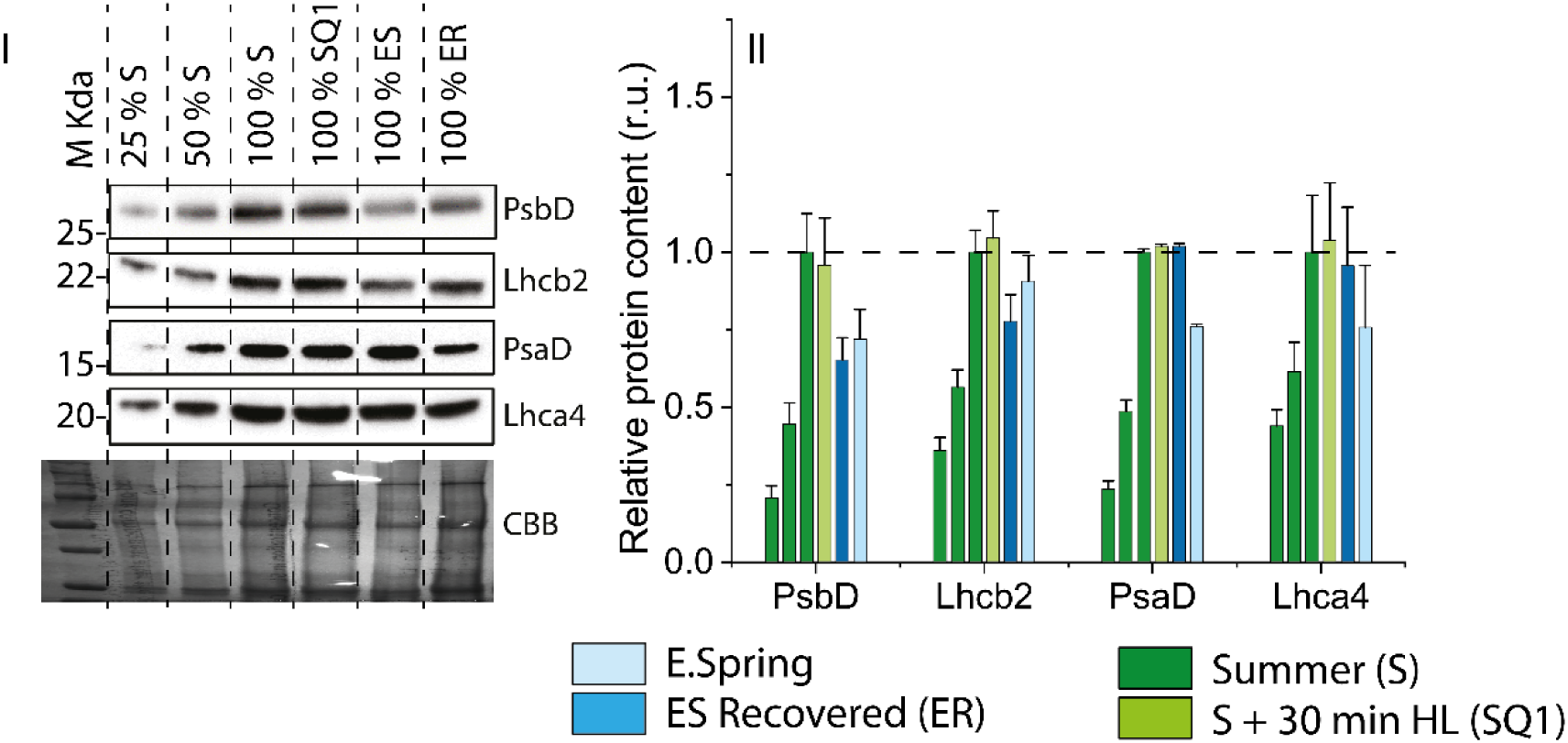

### Supplementary Information 6.

**Table 1.**
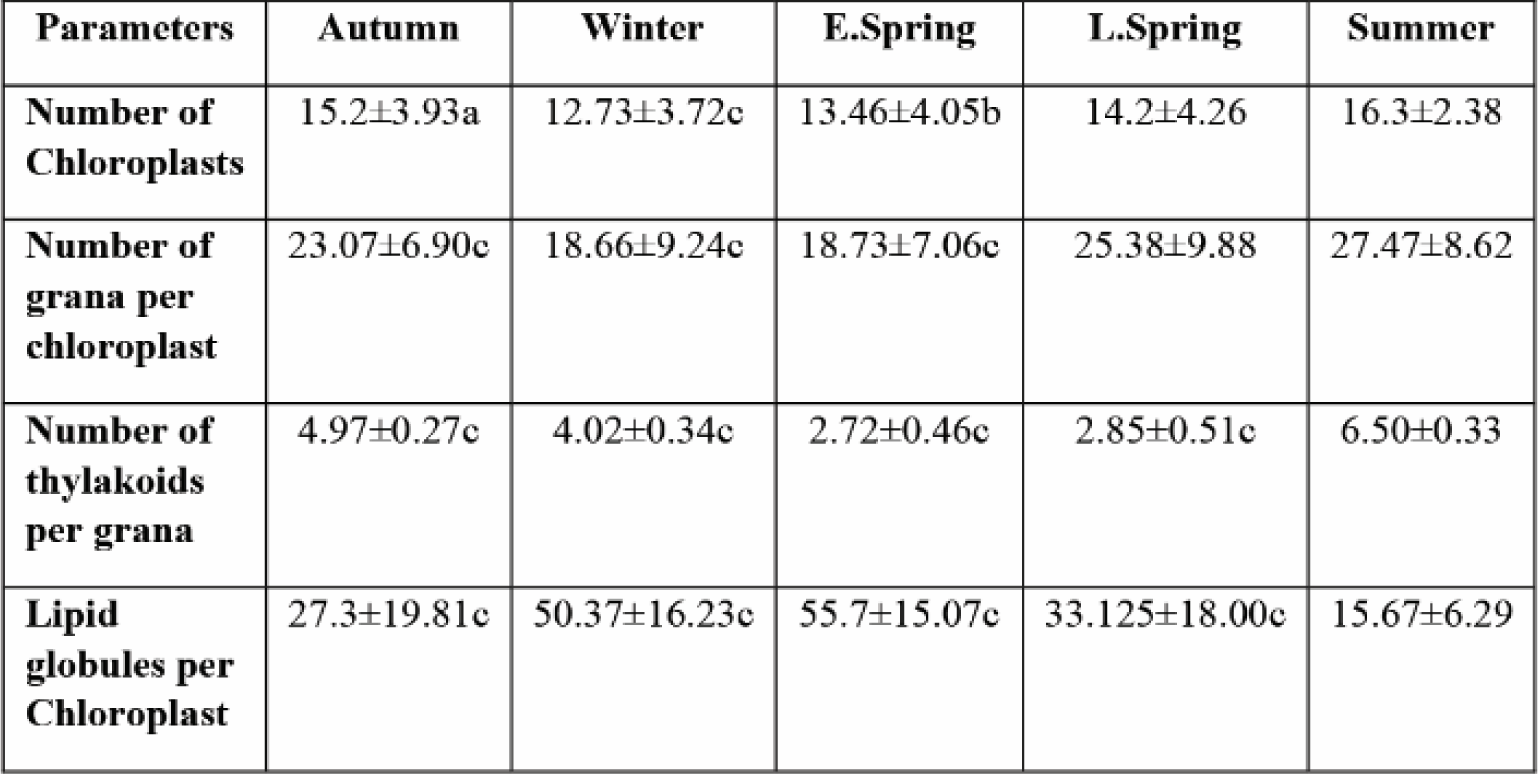
Quantitative analysis of seasonal changes in chloroplast ultrastructure as seen in Transmission electron microscopy. Statistical significance levels are referred as a, b, c denoting 99.95%, 99.99% and 99.999% level of significance.

**Table 2.**
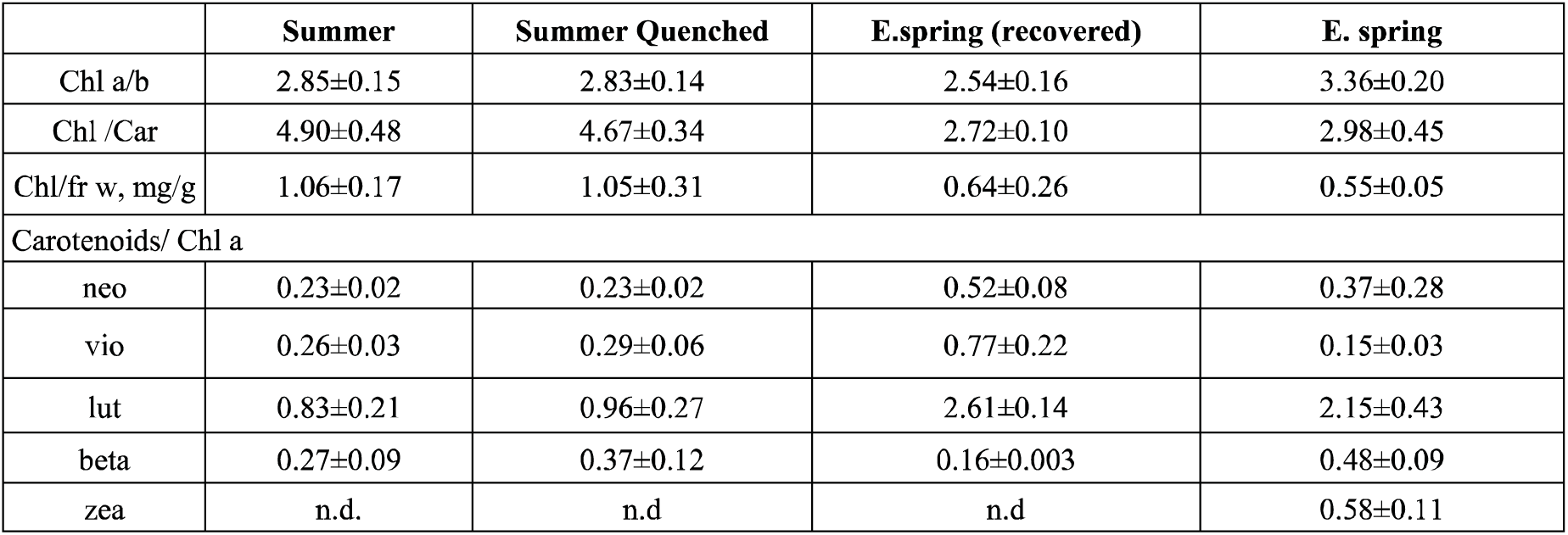
Pigment composition analysis by HPLC. Chl, Chlorophyll; fr w, fresh weight; neo, neoxanthin; vio violaxanthin; lut, lutein; beta, beta-carotene; zea, zeaxanthin. Shown is ±SD. n=3

**Table 3.**
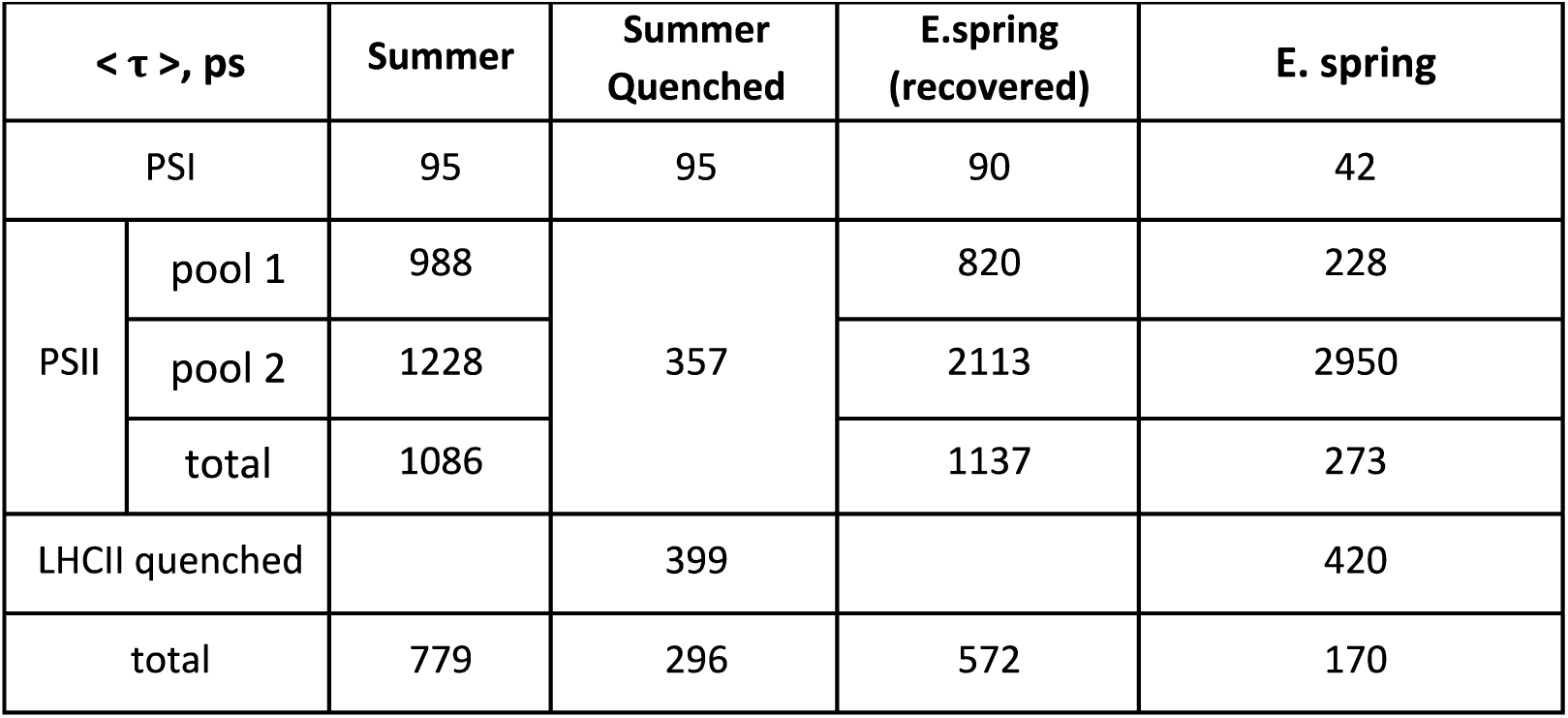
To assess differences in excited-state energy relaxation of different decaying components we calculated the average excited state relaxation time as < τ >= Σ *A*_*i*_ *Σ*_*i*_, where A_i_ are the relative areas of each Decay-associated spectra (DAS). DAS were obtained from global target analysis (Fig. 3).

**Table 4.** Percentages of total energy flow into PSII and PSI as calculated from Fig. 4.3 SI. For S state both PSII pools were taken into calculation

| Sample condition | PSI (CS and quenched) | PSII (CS) (Both pool taken together) | Comments |
| --- | --- | --- | --- |
| Summer unquenched | 50% | 30% | figure shows only main pool contribution |
| Summer quenched | 67% | 7.1% | detached and quenched LHCII not considered |
| E.Spring relaxed | 27% | 55% | small amount of photoinhibited PSII pool not considered |
| E.Spring quenched | 89% | 1.5% | small unquenched PSII pool and small quenched detached LHCII component not considered. |

